# LoRTIA Plus: a chemistry-agnostic, feature-first software package for long-read transcriptome annotation

**DOI:** 10.64898/2026.04.03.716279

**Authors:** Gábor Torma, Zsolt Balázs, Ádám Fülöp, Dóra Tombácz, Zsolt Boldogkői

**Affiliations:** Department of Medical Biology, Albert Szent-Györgyi Medical School, University of Szeged, Szeged, Hungary; Department of Quantitative Biomedicine, University of Zurich; Department of Medical Oncology and Hematology, University Hospital Zurich, Zurich, Switzerland

## Abstract

Long-read RNA sequencing (lrRNA-seq) enables direct reconstruction of full-length transcripts, yet existing annotation tools show variable performance across genomes and library chemistries, particularly for novel isoforms. We present LoRTIA Plus, a chemistry-agnostic suite for transcriptome annotation and reconstruction from lrRNA-seq data. LoRTIA Plus first detects and filters transcription start sites (TSSs), transcription end sites (TESs), and introns using adapter-aware and quality-based criteria, and evaluates read support before assembling high-confidence transcript models. We benchmarked LoRTIA Plus against bambu, FLAIR, IsoQuant, and NAGATA on KSHV transcriptomes with dense overlap, using a validated literature-supported boundary set, and on transcriptomes from three human cell lines from the Long Read RNA-seq Genome Annotation Assessment Project (LRGASP) sequenced with five long-read chemistries. On KSHV, LoRTIA Plus achieved the highest F1 scores for TSSs, TESs, and transcripts in both direct-cDNA and direct-RNA datasets by improving recall without sacrificing precision. Across human datasets, LoRTIA Plus consistently ranked among the top boundary annotators across all chemistries and was the best-performing tool in PCR-based libraries, while remaining highly competitive on native RNA. Junction- and isoform-level analyses show that LoRTIA Plus yields a rich, reproducible repertoire of novel isoforms and transcript boundaries from viral to human transcriptomes.

## Introduction

Accurate transcriptome annotation is fundamental to understanding gene expression, isoform diversity and regulatory complexity (1). A single gene locus can give rise to multiple isoforms via alternative RNA processing—including splicing, promoter usage, and polyadenylation—mechanisms that contribute to tissue specificity, development, and disease phenotypes (2–4). Inaccurate or incomplete annotation distorts isoform-level analyses and can compromise interpretation in both basic and translational studies (5). Long-read RNA sequencing (lrRNA-seq), as implemented by Pacific Biosciences (PacBio) and Oxford Nanopore Technologies (ONT), enables characterization of full-length mRNAs, non-coding RNAs (ncRNAs), and their isoforms, thereby reducing the need for assembly and improving the detection of complex patterns such as intron retention, recursive splicing, and alternative use of transcription start sites (TSSs) and transcription end sites (TESs) (6–8). Nevertheless, precise, reliable annotation remains challenging due to platform-specific biases, variable read accuracies and complex transcript architectures (9, 10).

Current lrRNA-seq analysis frameworks typically combine splice-aware aligners such as minimap2 (11) with reference- or genome-guided transcript reconstruction and quantification tools. The transcriptome annotator FLAIR (6) uses long reads aligned to a reference genome together with gene annotations and, optionally, short-read evidence to correct splice sites, collapse redundant isoforms and perform isoform-level analyses. IsoQuant (12) is a reference genome-based annotator that can operate either in a reference-guided mode, using existing gene annotations, or in a *de novo*, annotation-free mode, reconstructing and quantifying transcript models at exon and isoform resolution. Bambu (13) is also a reference-guided framework that takes aligned reads and genome annotations as input and uses a transcript-graph and machine-learning-based approach for context-aware quantification and discovery of novel isoforms relative to the reference. NAGATA (14) is specifically designed for compact viral genomes, integrating Nanopore-specific features to improve transcript boundary detection in densely packed transcriptomes. Many of these tools rely on prior annotations or are tuned to specific model systems, limiting discovery of novel isoforms. Conservative calling strategies can overlook biologically relevant variants, whereas permissive ones inflate false positives (9, 13–16).

To address these challenges, we developed LoRTIA Plus, a chemistry-agnostic, feature-first transcriptome analysis suite that extends our original LoRTIA (Long-read RNA-seq Transcript Isoform Annotator) framework, used in several viral transcriptomics studies (17–21) and here released for the first time as a standalone tool. LoRTIA Plus follows a feature-first strategy: it detects and filters TSSs, TESs and introns using adapter-aware and motif-based criteria where applicable, and statistically evaluates read support using Poisson modeling with Bonferroni correction for multiple testing, before reconstructing transcript models and quantifying isoforms across current ONT and PacBio chemistries. An additional adapter-independent mode enables consistent processing of ONT direct RNA sequencing (dRNA-seq) datasets.

Beyond compact viral testbeds, robust annotation must scale to complex human transcriptomes and diverse library chemistries. Community benchmarks such as the Long Read RNA-seq Genome Annotation Assessment Project (LRGASP) span multiple human cell lines and long-read protocols, including ONT dRNA-seq, cDNA-seq, CapTrap, and PacBio-based libraries (22). These benchmarks highlight recurring pitfalls: chemistry-dependent coverage and error profiles, truncated or chimeric molecules, inconsistent 5′/3′ end recovery, and inflation of spurious “novel” calls in permissive pipelines (22). Using the SQANTI3 framework for primary and extended structural classes (15, 16, 22), these datasets stress-test an annotator’s ability to recover known models while discovering biologically plausible isoforms without increasing the false positive rate. LoRTIA Plus implements a feature-first strategy that validates TSSs, TESs and introns before model assembly, using adapter- and motif-based cues together with statistical evaluation of read support to mitigate platform- and library-specific artifacts.

In this study, we benchmarked LoRTIA Plus against bambu (13), FLAIR (6), IsoQuant (12) and NAGATA (14) on the compact Kaposi’s sarcoma-associated herpesvirus (KSHV) transcriptome (20) that features densely overlapping transcription. We used a high-confidence reference set of promoter-proximal TSSs derived from Cap Analysis of Gene Expression (CAGE) and RNA Annotation and Mapping of Promoters for Analysis of Gene Expression (RAMPAGE) data, TESs supported by ONT dRNA poly(A) tails, and introns compiled from previously reported splice junctions (20, 23–25). In addition, we evaluated these annotators on LRGASP transcriptomes from three human cell lines (H1-hES, H1-DE and WTC11 iPSC) across five long-read chemistries and two sequencing platforms. In the following sections, we quantify precision, recall, F1 scores, category-level recovery and novel isoform discovery across chemistries and annotators.

## Materials and Methods

### Overview of the LoRTIA Plus pipeline

The LoRTIA Plus framework is a three-stage workflow that progressively filters and validates long-read RNA-sequencing data to produce high-resolution, biologically reliable transcript annotations. Unlike conventional annotators, LoRTIA Plus first applies stringent quality control and statistical validation to candidate transcriptional features (TSSs, TESs, introns), and only then assembles them into full-length transcript isoforms. To ensure completeness, the software re-screens the BAM file to verify that every reconstructed transcript is supported by valid reads. This filter-first, assemble-later strategy differs fundamentally from assembly-first approaches and aims to reduce artifacts while maximizing endpoint accuracy.

### Step 1 — Adapter-aware QC and read tagging

LoRTIA Plus iterates over every read in a BAM/SAM file and aligns the 5′ and 3′ terminal sequences to the expected adapter sequences using the Smith–Waterman algorithm. Reads are flagged as adapter-positive only if the alignment score meets or exceeds a minimum threshold, and only reads carrying both adapters are retained. In cDNA-seq libraries, LoRTIA Plus uses adapter detection to determine transcript strand orientation. Internal priming and template switching can occur at stretches of consecutive adenines, producing erroneous 3′ ends; therefore, 3′ ends within homopolymer A runs are flagged (17). In addition, introns preceded by long insertions—often arising from chimeric molecules—are removed. Adapter-aware filtering is critical; the strand-switching oligo used during library preparation has a defined 5′ sequence, including a 3G tail that anneals to the non-templated cytosines added by the reverse transcriptase; without explicitly matching this adapter, 5′ completeness cannot be reliably assessed. For transparency, LoRTIA Plus outputs tab-delimited QC logs enumerating (i) invalid or missing 5′ adapter signatures, (ii) 3′ end candidate sites consistent with internal priming, and (iii) introns suspected to arise from template switching.

### Step 2 — Statistical validation of transcript features

Candidate TSSs and TESs are identified as local maxima in read-start or read-end distributions; only sites supported by ≥2 reads and exceeding a defined coverage ratio are retained. Features within ±10 nt are collapsed into clusters, and significance is evaluated using a Poisson distribution, where *k* is the read-end count at the cluster peak and λ is the average read-end count within a ±50-nt window. P-values are adjusted for multiple hypothesis testing using the Bonferroni method by multiplying the nominal P-values by the number of evaluated features. In cDNA libraries, TESs are further filtered for template-switching artifacts (17). This procedure re-evaluates candidate poly(A) sites from LoRTIA outputs and classifies them as genuine TESs or potential template-switching artifacts based on read-support patterns, specifically the proportion of reads ending at a given TES and the number of consecutive adenines at that TES. This additional filtering reduces spurious TES calls while preserving well-supported transcript-end boundaries. For introns, additional filters remove rare splice junctions located within 15 nt of frequent junctions in order to reduce sequencing error-induced artifacts; canonical donor–acceptor splice-site motifs (GT/AG, GC/AG or AT/AC) are strictly required. Short homology sequences (SHSs)—direct repeats of ≥3 nt at donor–acceptor boundaries—are flagged to identify potential template-switching artifacts and remove erroneous junctions. This multilayered validation yields a high-confidence set of TSSs, TESs and introns.

### Step 3 — Transcript annotation

Validated features are integrated into full-length isoforms using strictly filtered reads. Only reads initiating within ±10 nt of an accepted TSS and terminating within ±10 nt of a validated TES are retained; introns must correspond to validated donor–acceptor pairs with canonical motifs. Reads containing non-canonical introns or large unsupported deletions are excluded. From this strictly filtered read set, LoRTIA Plus creates transcript models and reports them in GFF3 format, while also outputting an annotated BAM file for debugging purposes or quantitative analyses.

### KSHV benchmark datasets

For benchmarking, we analysed KSHV transcriptomes generated with Oxford Nanopore Technologies (ONT) long-read sequencing. The datasets included dRNA-seq (kit dRNA-002) and direct cDNA sequencing (kit DCS-109) from Shekhar et al. (23) (SRA BioProject PRJNA975091), as well as additional dcDNA-seq runs (DCS-109) from Prazsák et al. (20) (ENA PRJEB60022). To obtain an independent reference for transcript boundaries, we integrated previously published KSHV annotations. Promoter-proximal TSSs were taken from the KSHV 2.0 and KSHV 3.0 annotations and from Shekhar et al. and Ye et al., and were retained only if they had been explicitly reported as TSSs and were supported by CAGE and/or RAMPAGE peaks (20, 23–25). TESs were taken from KSHV 2.0/3.0 and from Shekhar et al., and were retained only when they coincided with TES positions supported by ONT dRNA 3′-end [poly(A)] signals in either our datasets or those of Shekhar et al. Introns were compiled from splice junctions reported across KSHV 2.0, KSHV 3.0, Shekhar et al. and RAMPAGE-based studies. This literature-derived, assay-supported set of TSSs, TESs and introns served as a high-confidence reference for benchmarking LoRTIA Plus and the alternative tools. All accession numbers and dataset details are listed in **Supplementary Table S1A**.

### Human benchmark datasets

We also evaluated transcript annotation tools using the community-driven LRGASP study (22), including three human cell types: WTC11 induced pluripotent stem cells (iPSCs), H1 human embryonic stem cells (H1-hES) and H1-derived definitive endoderm (H1-DE). For each cell type, libraries were generated using five long-read chemistries: ONT dRNA-seq, ONT dcDNA-seq, ONT CapTrap, PacBio cDNA and PacBio CapTrap. Long-read data for all experiments were obtained from the ENCODE Portal. This design provides a diverse and well-controlled benchmark for assessing isoform recovery and de novo transcript reconstruction across chemistries while keeping the underlying biology fixed. Per-experiment accessions (ENCSR/ENCFF) are listed in **Supplementary Table S1B** (see also the Data Availability section).

### Reference resources and long-read alignment

We used the human reference genome GRCh38.p14 with gene models from GENCODE Release 49 (GTF) (26). For KSHV, the reference genome was the GQ994935.1 GenBank assembly. Long-read RNA-seq data were aligned with minimap2 v2.30 using spliced presets appropriate for the library type: -x splice for ONT/PacBio cDNA and CapTrap libraries (reads were aligned in an unstranded manner; LoRTIA Plus subsequently assigns strand using adapter signatures where applicable), and -x splice -uf -k14 for ONT dRNA sequencing (strand-aware, noisy reads) (11). SAM/BAM conversion, sorting and indexing were performed with SAMtools v1.22.1 (27).

### Evaluation metrics

Human TSS annotation accuracy was evaluated using refTSS (FANTOM/FANTOM5 promoters) as an independent reference (28). In refTSS, each entry represents a promoter-associated TSS cluster derived from CAGE data and represented as a genomic interval rather than a single nucleotide; accordingly, each refTSS interval was treated as one promoter unit. Because ONT dRNA-seq was not included in TSS benchmarking, concordance between predicted TSSs and reference promoter units was assessed for the remaining chemistries using a strand-aware ±10-nt tolerance. Within the reference set, a promoter unit was counted as a true positive (TP) for a given annotator if it was matched by at least one predicted TSS; unmatched active promoter units were counted as false negatives (FN), and predicted TSSs not matching any active promoter unit were counted as false positives (FP).

TES accuracy was evaluated using two independent poly(A)-site resources, PolyASite 2.0 (29) and PolyA_DB v4 (30). For PolyASite 2.0, reference TESs were used at their reported peak position, and predicted TESs were considered concordant if they fell on the same chromosome and strand within ±25 nt of a PolyASite peak. For PolyA_DB v4, TES concordance was also evaluated with a ±25-nt tolerance, consistent with the database construction strategy in which nearby cleavage events are clustered within approximately 25 nt. Only TESs recovered by at least one annotator in the corresponding sample were retained in the active reference set. Within this set, a reference TES was counted as a TP if matched by at least one predicted TES, unmatched TESs were counted as FN, and predicted TESs not matching any reference TES were counted as FP.

To evaluate end-to-end transcript boundary reconstruction, we defined reference transcript-end units as unique (promoter unit, TES site) pairs. For each sample, only those transcript-end units that were recovered by at least one annotator were retained in the reference set. A predicted transcript was accepted as correct at the transcript level only if it matched both ends jointly, that is, if its TSS mapped to the same refTSS promoter unit within ±10 nt and its TES matched the same reference TES site within the corresponding TES tolerance. TP, FP and FN were then computed on these unique transcript-end units.

For each annotator × chemistry × cell-line combination, we tallied TP, FP and FN separately for TSSs, TESs and transcript-end units and computed precision, recall and F1 as precision = TP/(TP+FP), recall = TP/(TP+FN) and F1 = 2·precision·recall/(precision+recall).

To restrict benchmarking to reference features that were relevant in a given sample, in both the human LRGASP and KSHV analyses, a reference feature was retained only if it was recovered by at least one annotator in the corresponding benchmark on the correct strand and within the relevant positional tolerance. These reference sets were used for all downstream TP/FP/FN calculations.

For the KSHV benchmarks, we applied the same general benchmarking framework. The viral reference consisted of literature- and experimentally supported KSHV features compiled from Shekhar et al., Prazsák et al. and Arias et al. Candidate TSSs, TESs and transcript-end features were retained in the KSHV reference only if at least one annotator recovered the corresponding position or feature within the defined positional tolerance. These reference sets were then used to score TSS-, TES- and transcript-level recovery in the viral benchmark.

### SQANTI3-based structural annotation of transcript isoforms

For human structural comparisons, transcript models produced by each annotator were classified against the GENCODE v49 reference annotation using SQANTI3 into the primary categories full-splice match (FSM), incomplete-splice match (ISM), novel in catalog (NIC) and novel not in catalog (NNC), together with the additional classes Antisense, Genic Intron, Genic Genomic, Intergenic and Fusion (15, 16). To complement the boundary-based benchmarking, we also assessed intron-level structural performance in the human LRGASP data by focusing on FSM and ISM transcript models, which reflect exact or partial recovery of reference intron chains. These structural summaries were interpreted within the same annotation space used for the boundary-level analyses, ensuring direct comparability between structural and end-based benchmarking results.

All SQANTI3-derived output files for the human analyses, including the GTF annotations and the corresponding classification.txt and junction.txt tables for each annotator × chemistry × cell-line combination, are available from FigShare via the link provided in the Data Availability section.

### Benchmarking configurations of transcript annotation tools

All transcript annotation tools were run on identical inputs, including the same reference sequences, alignment files and downstream comparison framework, using standardized settings based on publicly documented default workflows and author recommendations, so that observed differences primarily reflect method-specific behavior rather than arbitrary configuration choices. The benchmark configurations of the evaluated transcript annotation tools are summarized in **Table 1**. LoRTIA Plus was run using its standard feature-first, end-aware workflow, in which transcript features are first identified and filtered and then used for transcript reconstruction. For cDNA-based datasets, chemistry-appropriate adapter settings were applied, whereas Oxford Nanopore direct RNA datasets were processed in direct RNA mode. bambu was run in de novo transcript discovery mode with NDR = 1 and single-exon discovery enabled. IsoQuant was run with platform-specific presets for Oxford Nanopore and PacBio data. NAGATA was run using its default parameter set. FLAIR was run using its standard workflow with correction followed by transcript collapse. SQANTI3 was used for structural classification of transcript models against the corresponding reference annotation, using GENCODE v49. For all tools, the same input alignments were used, and downstream outputs were processed within a unified benchmarking framework.

**Table 1.**
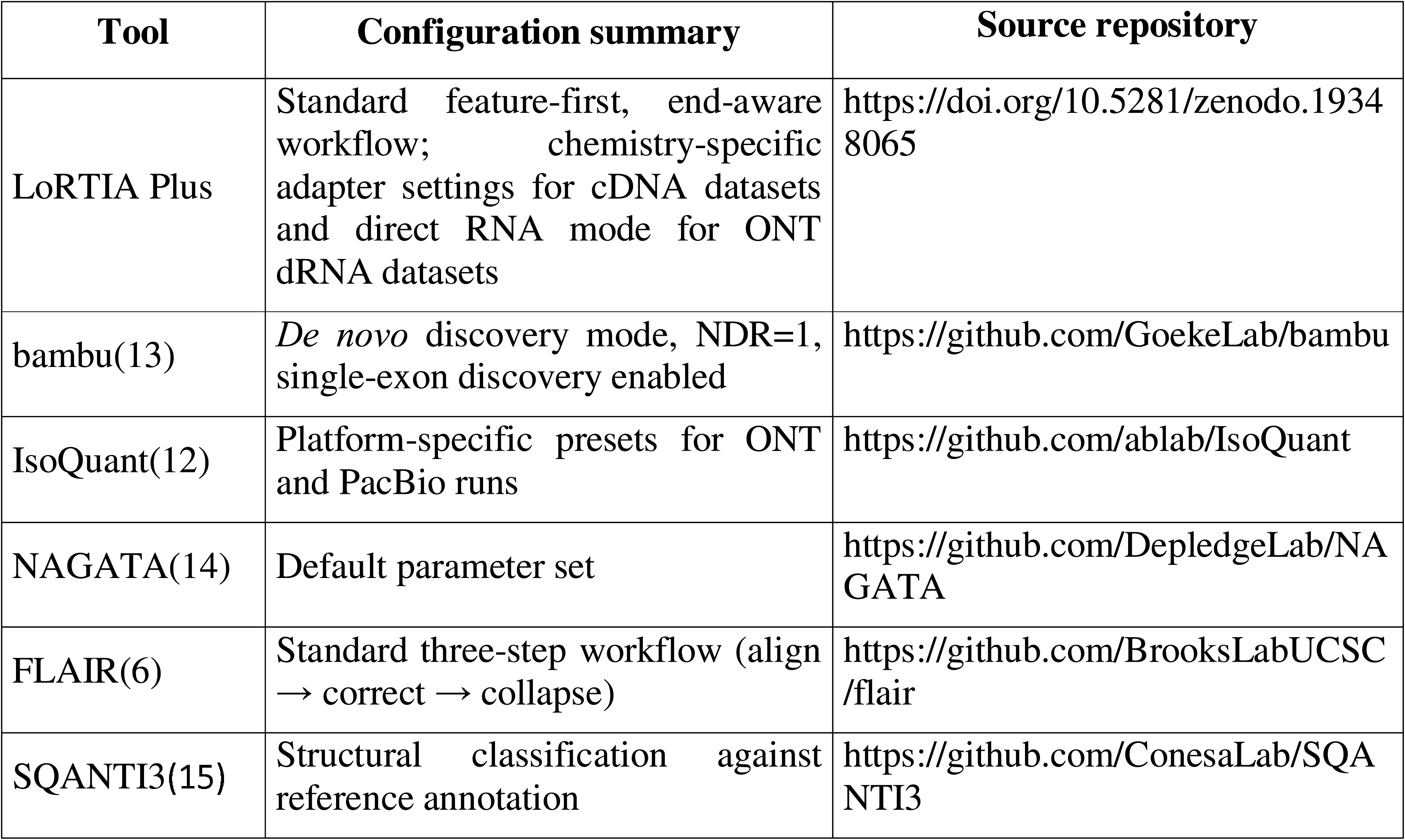
Overview of the transcript annotation tools and their execution parameters.

## Results

### ONT Nanopore chemistry benchmarking on the KSHV transcriptome

To establish an independent, biologically supported reference for transcript-boundary detection, we compiled a high-confidence set of KSHV transcript start sites (TSSs) and transcript end sites (TESs) from orthogonal evidence sources (see Methods). Validated TSSs were restricted to positions supported by published CAGE and RAMPAGE datasets, whereas validated TESs were restricted to 3′ ends supported by poly(A)-anchored ONT direct RNA reads. For each annotator–chemistry combination, recovery of validated sites was quantified within a permissive positional tolerance (see Methods), and precision, recall and F1 were calculated from TP, FP and FN counts **(Figure 1A; Supplementary Table S2A)**.

**Figure 1.**
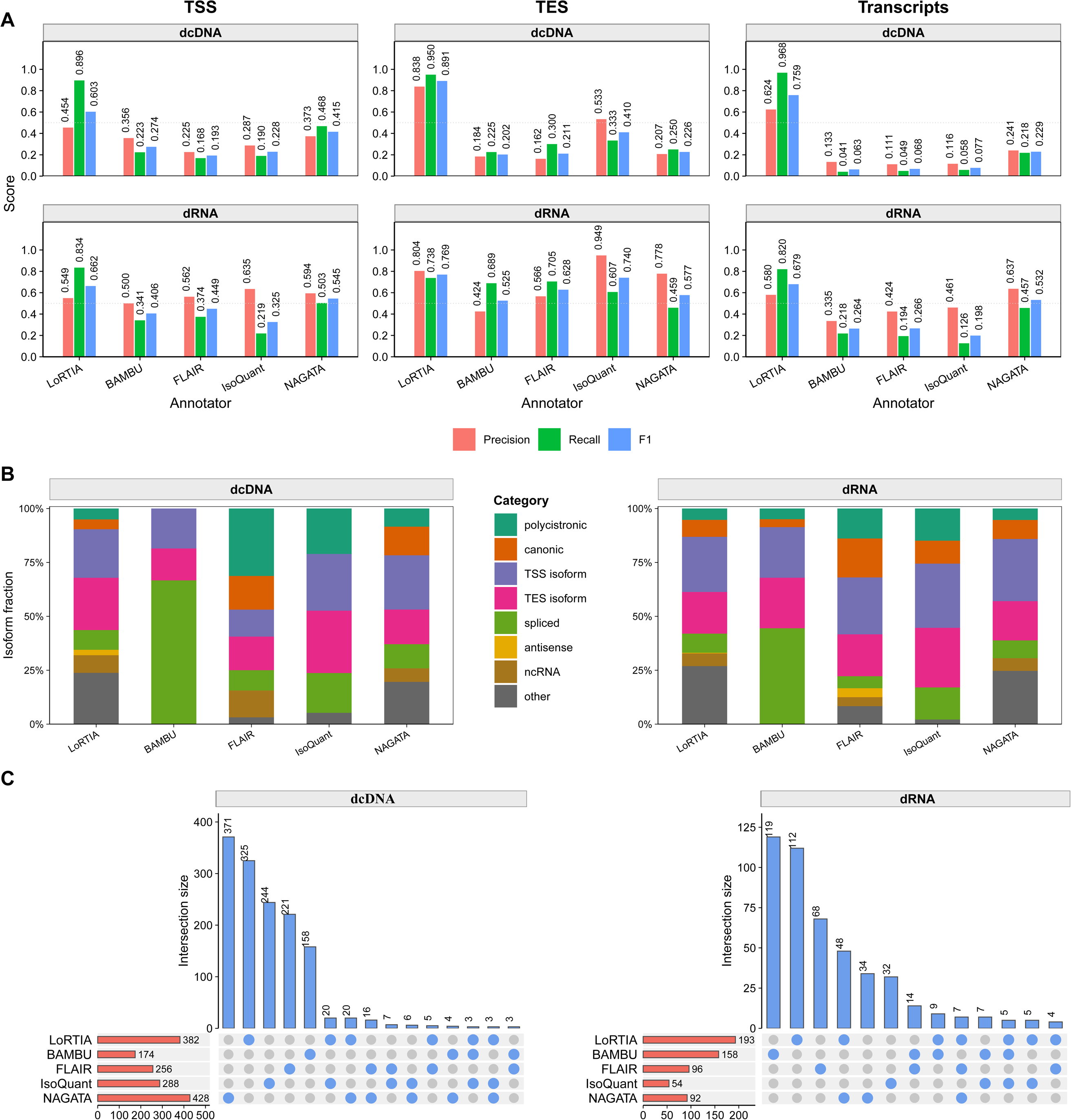
Benchmarking of KSHV transcript annotation across ONT direct cDNA and direct RNA chemistries. **(A)** Precision, recall and F1 values for TSS, TES and transcript-level recovery across five annotators (LoRTIA Plus, bambu, FLAIR, IsoQuant and NAGATA) in direct cDNA (dcDNA) and direct RNA (dRNA) datasets. Scores were calculated from TP, FP and FN counts using the biologically supported KSHV reference set defined in Methods. **(B)** Transcript-class composition of reference-matched transcript models recovered by each annotator in dcDNA and dRNA datasets. Detailed transcript classes were recoded into broader categories (polycistronic, canonic, TSS isoform, TES isoform, spliced, antisense, ncRNA and other), and bars show the relative fraction of each category within the matched transcript set of each tool. **(C)** UpSet-style summary of unmatched (false-positive) transcript models in dcDNA and dRNA datasets. Horizontal bars indicate the total number of unmatched transcript models produced by each annotator, top bars show the sizes of retained intersections among annotator-specific false-positive sets, and the dot matrix indicates the annotator combinations defining each intersection. For clarity, only intersections of size >2 are shown. Alt text: Benchmark figure comparing five annotation tools across KSHV ONT direct cDNA and direct RNA datasets, including boundary-detection accuracy, matched transcript classes, and false-positive overlap patterns.

For TSS detection, LoRTIA Plus achieved the highest F1 under both ONT chemistries. In direct cDNA, it reached a TSS F1 of 0.603, outperforming all competing annotators **(Figure 1A; Supplementary Table S2A)**. This advantage was driven primarily by substantially higher recall (0.896) than that of the other methods (0.168–0.468), while also retaining the highest precision in the benchmark (0.454). A similar pattern was observed for direct RNA, where LoRTIA Plus again produced the best TSS F1 (0.662), owing mainly to improved recall (0.834), with precision (0.549) remaining within the range of competing methods. Collectively, these results show that LoRTIA Plus most effectively recovered validated KSHV TSSs across both ONT chemistries.

For TES detection, the clearest separation between methods was observed in direct cDNA. Here, LoRTIA Plus achieved a TES F1 of 0.891, substantially exceeding all competing annotators **(Figure 1A; Supplementary Table S2A)**. This advantage reflected improvements in both performance components, as LoRTIA Plus combined the highest precision (0.838) with the highest recall (0.950), whereas all other tools performed markedly worse on both axes. In direct RNA, differences between methods were smaller, but LoRTIA Plus still achieved the highest TES F1 (0.769). The closest competitor was IsoQuant, which reached very high precision (0.949) but lower recall (0.607), whereas LoRTIA Plus maintained a more balanced precision–recall profile (0.804/0.738), resulting in the best overall F1. Thus, LoRTIA Plus ranked first for TES identification under both ONT chemistries in this biologically validated KSHV benchmark.

Because accurate transcript reconstruction depends on correct pairing of transcript boundaries, we next evaluated performance at the transcript level. Consistent with the boundary-level results, LoRTIA Plus provided the most accurate end-to-end transcript reconstruction under both ONT chemistries. In direct cDNA, it achieved a transcript-level F1 of 0.759, clearly outperforming all competing annotators (**Figure 1A; Supplementary Table S2B**). This reflected both high precision (0.624) and near-complete recall of validated transcript models (0.968), whereas competing methods recovered only a small fraction of reference transcripts (recall = 0.041–0.218). In direct RNA, LoRTIA Plus again yielded the highest transcript-level F1 (0.679). The closest competitor was NAGATA (F1 = 0.532), which showed slightly higher precision (0.637) but substantially lower recall (0.457) than LoRTIA Plus (precision = 0.580, recall = 0.820). These findings indicate that the boundary-detection advantage of LoRTIA Plus translates directly into improved full-length transcript reconstruction on the KSHV transcriptome.

Finally, we asked whether these performance differences were accompanied by systematic shifts in transcript-model composition across annotators **(Figure 1B)**. To this end, detailed transcript classes were recoded into broader categories (polycistronic, canonic, TSS isoform, TES isoform, spliced, antisense, ncRNA and other) using a rule-based scheme, and the relative contribution of each category was summarized for the matched transcript sets recovered by each tool **(Supplementary Table S2C, D)**. The resulting profiles were clearly annotator-specific and broadly stable across chemistries: LoRTIA Plus and NAGATA yielded the most compositionally diverse matched sets, bambu was dominated by spliced matches, FLAIR showed comparatively greater representation of polycistronic models, whereas IsoQuant was relatively enriched for transcript-isoform classes, particularly TSS and TES isoforms. These results indicate that annotator choice affected not only recovery of validated transcript boundaries, but also the category balance of the transcript models retained in the benchmark.

To further resolve annotator-specific output patterns, we summarized the composition of unmatched transcript models in **Supplementary Table S2E, F** and their overlap structure in **Figure 1C**. False-positive burdens were consistently higher in direct cDNA than in direct RNA, and in both chemistries most unmatched transcript models were annotator-specific rather than broadly shared across methods. In direct cDNA, the largest false-positive sets were produced by NAGATA and LoRTIA Plus, whereas in direct RNA unmatched calls were fewer overall but still showed clear annotator-specific structure, with an additional shared subset between LoRTIA Plus and NAGATA. Together, these results show that annotator differences are driven as much by method-specific patterns in how false-positives are generated as by sensitivity to validated transcript boundaries.

### Global structural properties of human long-read transcriptomes

We evaluated the above-mentioned transcript annotation tools on long-read sequencing (LRS) data from three human LRGASP cell lines—H1 human embryonic stem cells (H1-hES), definitive endoderm derived from H1 (H1-DE), and WTC11 iPSCs—generated using five long-read library chemistries (ONT-dRNA, ONT-cDNA, ONT-CapTrap, PacBio cDNA, PacBio-CapTrap). LoRTIA Plus was compared with bambu, IsoQuant, NAGATA and FLAIR using SQANTI3 primary structural categories—full-splice match (FSM), incomplete-splice match (ISM), novel in catalog (NIC) and novel not in catalog (NNC)—together with additional classes (antisense, genic, genic_intron, intergenic, fusion). Although NAGATA is optimized for compact viral genomes, we included it here for completeness and for consistent cross-tool comparisons. Cell line–aggregated summaries of SQANTI3 class composition, transcript length distributions, coding/non-coding balance and gene-level isoform complexity are shown in **Figure 2A–D**, while per-cell-line details are provided in **Supplementary Figure S1A-D**. The corresponding numerical summaries are reported in **Supplementary Tables S3A–S3C**.

**Figure 2.**
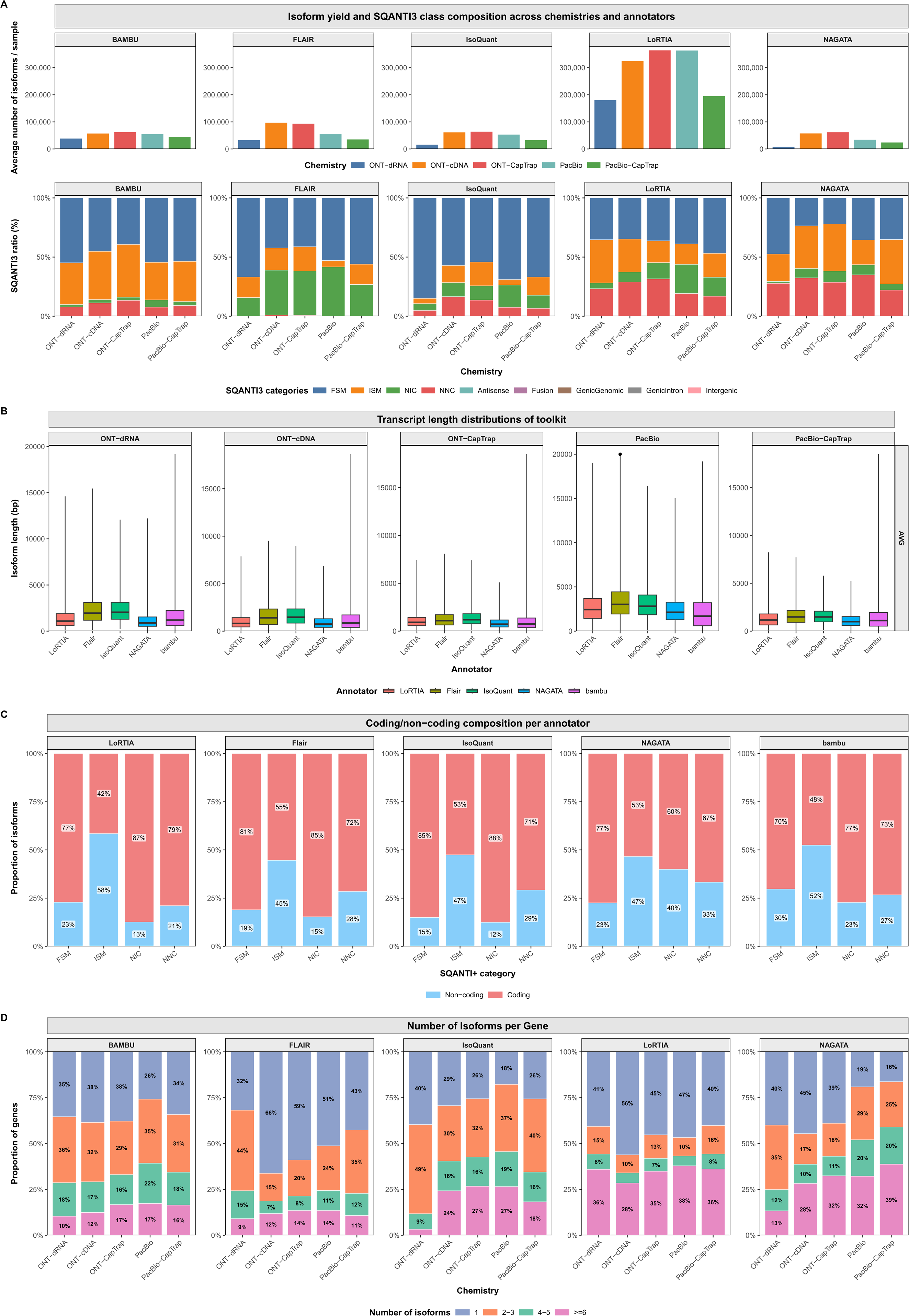
Cross-chemistry benchmarking of isoform annotation across three human LRGASP cell lines. Comparison of five long-read chemistries (ONT-dRNA, ONT-cDNA, ONT-CapTrap, PacBio cDNA, PacBio-CapTrap) across three cell lines (H1-hES, H1-DE, WTC11) and five annotators (LoRTIA Plus, bambu, FLAIR, IsoQuant, NAGATA). Main panels summarize cell line–averaged values across the three cell lines. **(A)** Mean isoform yield across the three cell lines, with stacked SQANTI3 class composition shown for each annotator–chemistry combination. **(B)** Cell line–averaged transcript-length distributions for each annotator and chemistry. **(C)** Mean coding versus non-coding composition across the three cell lines for each annotator and chemistry. **(D)** Mean gene-level isoform complexity across the three cell lines for each annotator and chemistry. Alt text: Cross-chemistry benchmark across three human LRGASP cell lines, summarizing isoform yield, transcript length, coding potential, and gene-level isoform complexity across annotators.

SQANTI3-based classification showed that, across chemistries and cell lines, the majority of isoforms fall into catalog-compatible FSM/ISM classes, but each annotator exhibits a distinct and highly reproducible category profile **(Figure 2A; Supplementary Table S3A)**. In this setting, IsoQuant displayed the most catalog-centered overall profile, whereas bambu was relatively enriched for ISM models, consistent with a reconstruction regime in which partially truncated isoforms remain splice-architecture–concordant with the reference. FLAIR and LoRTIA Plus both recovered substantial catalog-external transcript diversity, but with different emphases: FLAIR was skewed toward NIC, whereas LoRTIA Plus showed a broader novelty profile with a stronger NNC component. NAGATA yielded the most permissive overall category distribution, with comparatively large contributions from catalog-external and auxiliary classes. These patterns remained stable across the three tested cell lines, indicating that the primary SQANTI3 category distributions are predominantly shaped by annotator strategy rather than by the underlying biological system **(Figure 2A; Supplementary Figure S1A)**.

Transcript length distributions revealed additional chemistry- and tool-specific features **(Figure 2B; Supplementary Table S3B)**. Across tools, library chemistry was the dominant determinant of transcript length, with the longest pooled distributions observed in PacBio cDNA, intermediate distributions in ONT-dRNA and PacBio-CapTrap, and shorter distributions in ONT-CapTrap and ONT-cDNA **(Figure 2B; Supplementary Figure S1B; Supplementary Table S3B)**.

The balance between coding and non-coding transcripts depended on both library chemistry and annotator **(Figure 2C; Supplementary Table S3A)**. Across chemistries, coding fractions were highest in PacBio cDNA (∼0.836), intermediate and closely similar in ONT-dRNA and PacBio-CapTrap (∼0.692–0.693), and lower in ONT-CapTrap (∼0.586) and ONT-cDNA (∼0.557). Across annotators (aggregated over chemistries and cell lines), IsoQuant yielded the highest coding proportion (∼0.771), followed by LoRTIA Plus (∼0.683) and FLAIR (∼0.633), whereas bambu (∼0.586) and especially NAGATA (∼0.532) were lower. Together, these trends indicate that chemistry sets the overall coding/non-coding baseline, while annotator choice imposes a reproducible shift around that baseline **(Supplementary Figure S1C).**

Gene-level isoform complexity further separated the annotators into distinct groups **(Figure 2D; Supplementary Table S3C)**. FLAIR produced the most compact gene models (mean ∼2.81 isoforms per gene; fraction of genes with ≥6 isoforms ∼12.2%), whereas bambu (∼3.30; ≥6 ∼14.8%) and IsoQuant (∼3.87; ≥6 ∼21.8%) were intermediate. LoRTIA Plus and NAGATA yielded the richest isoform repertoires per gene (LoRTIA Plus ∼11.44 and NAGATA ∼5.49 isoforms per gene), with the highest fractions of genes harboring ≥6 isoforms (∼34.3% for LoRTIA Plus; ∼30.5% for NAGATA), consistent with their broader contribution to catalog-external transcript diversity. These patterns were reproducible across the three cell lines **(Supplementary Figure S1D)**.

Junction-level support patterns further indicated that annotators differed not only in the amount of catalog-external splice novelty they reported, but also in its reproducibility across tools and chemistries. Because these patterns are most directly relevant to transcript novelty outside the reference catalog, they are examined separately in the following section.

### Transcript boundary detection across annotators and long-read chemistries

Using the same three human LRGASP cell lines and five long-read library chemistries described above, we next compared transcript boundary detection (TSS and TES) across five annotators (LoRTIA Plus, bambu, FLAIR, NAGATA and IsoQuant). For each annotator–chemistry–cell-line combination we computed boundary-level precision, recall and F1 separately for TSSs and TESs. Differences between annotators were assessed per chemistry using an ANOVA with cell line included as a blocking factor, followed by pre-specified cell-line–paired t-tests comparing LoRTIA Plus versus each competing annotator with Benjamini–Hochberg (BH) FDR correction (see Methods). Given n=3 cell lines, we emphasize effect sizes alongside P-values. TSS benchmarking was restricted to the four PCR-based chemistries, because ONT-dRNA libraries frequently exhibit 5′-end truncation that compromises precise TSS localization; TES benchmarking included all five chemistries including ONT-dRNA. Detailed metric values are provided in **Supplementary Table S4A** (TSS) and **Supplementary Table S5A** (TES). Full ANOVA outputs and paired comparisons are reported in **Supplementary Tables S4B–S4C** (TSS) and **S5B–S5C** (TES), and summarized in **Figure 3A**, **C**, **E** for TSS and **Figure 3B**, **D**, **E** for TES.

**Figure 3.**
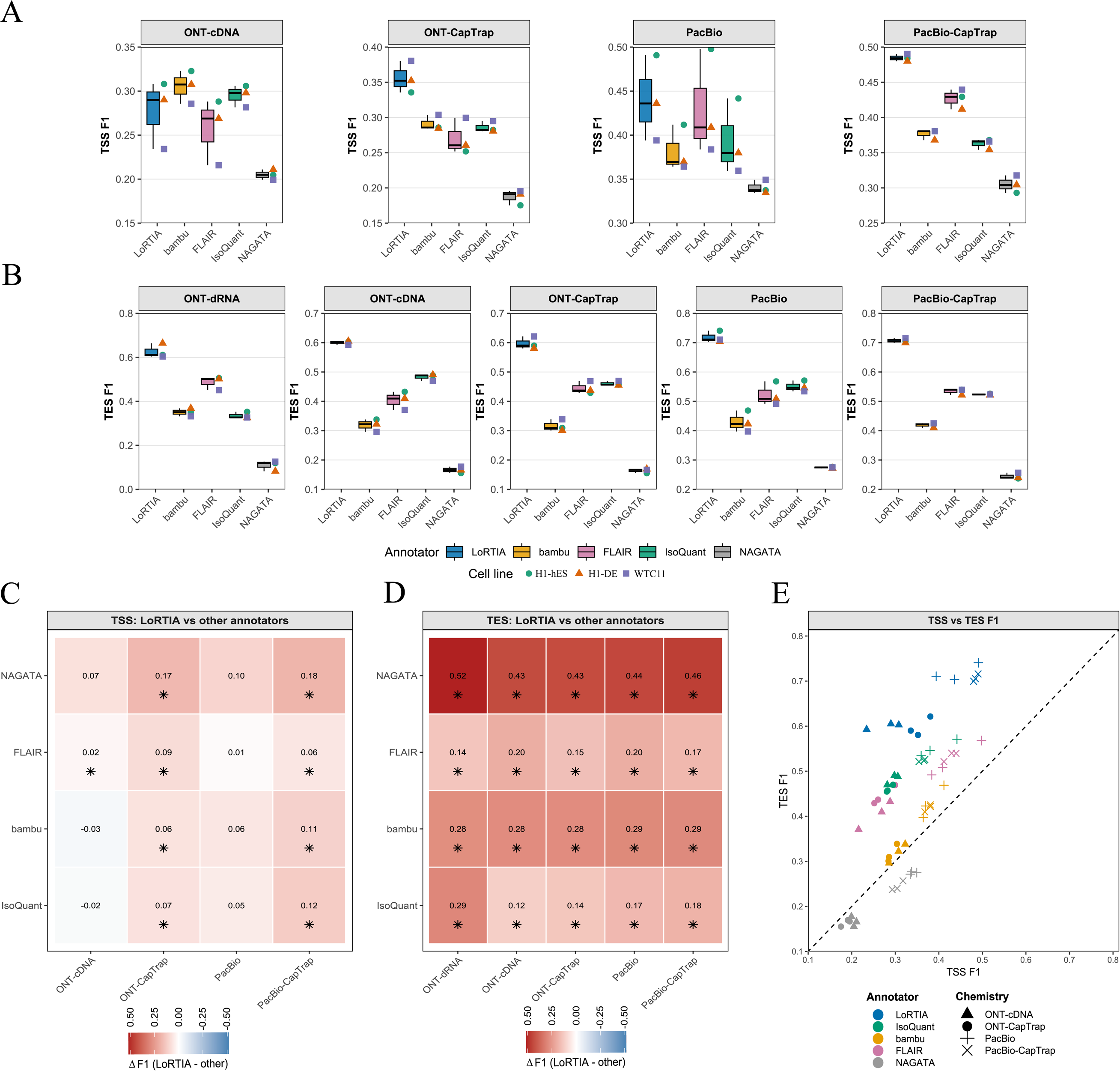
Chemistry- and annotator-dependent performance of TSS and TES detection across long-read transcriptome annotation tools. F1 scores (harmonic mean of precision and recall for recovery of benchmark-defined transcription start sites (TSSs) and transcription end sites (TESs) within the allowed positional tolerance; see Methods) were calculated for each combination of cell line (H1-hES, H1-DE and WTC11), long-read sequencing chemistry and annotator (LoRTIA Plus, bambu, FLAIR, NAGATA and IsoQuant). TSS was evaluated only for the four PCR-based chemistries, because ONT-dRNA libraries frequently exhibit 5′-end truncation that compromises precise TSS calling, whereas TES was evaluated for all five chemistries, including ONT-dRNA. For each chemistry, annotator effects were tested by ANOVA with cell line as a blocking factor, followed by pre-specified paired t-tests comparing LoRTIA Plus with each competing annotator across cell lines and Benjamini–Hochberg correction for multiple testing. **(A, B)** Boxplots of TSS **(A)** and TES **(B)** F1 scores across chemistries. Within each chemistry, the x-axis shows the annotators, boxplots summarize the distribution across the three cell lines, and overlaid points indicate individual cell-line values. These panels show how 5′- and 3′-boundary detection accuracy varies across annotators and library chemistries. **(C, D)** Heatmaps of pairwise F1 differences between LoRTIA Plus and each alternative annotator for TSS **(C)** and TES **(D)**. Each tile represents one chemistry–annotator comparison and reports the mean difference across cell lines, calculated as ΔF1 = F1(LoRTIA Plus) − F1(other annotator). Values are shown numerically and by a diverging color scale, with positive values indicating higher F1 for LoRTIA Plus. Asterisks denote comparisons significant after Benjamini–Hochberg correction (FDR < 0.05) based on paired t-tests across the three matched cell lines; significance therefore reflects both the magnitude and the cross-cell-line consistency of ΔF1, not effect size alone. **(E)** Scatter plot comparing TSS and TES F1 values for each cell line × chemistry × annotator combination. Points are colored by annotator and shaped by sequencing chemistry. The dashed diagonal (y = x) indicates equal TSS and TES performance; points above the line indicate stronger TES than TSS detection, whereas points below the line indicate the opposite. This panel provides an integrated view of how each annotator balances 5′- and 3′-boundary accuracy across long-read chemistries. Alt text: Comparison of TSS and TES detection performance across annotators and long-read chemistries, including F1 distributions, pairwise differences relative to LoRTIA Plus, and joint TSS–TES performance.

Across library preparations, annotator choice produced reproducible accuracy shifts that were largely concordant across the three cell lines. To summarize the practical magnitude of these differences, we report chemistry-resolved effect sizes as ΔF1 = F1(LoRTIA Plus) − F1(other), averaged across cell lines. The resulting ΔF1 matrices are shown as heatmaps **(Figure 3C–D)**, while absolute F1 distributions are shown as box/point plots **(Figure 3A–B and 3E)**. Overall, annotator-dependent separation was more pronounced and more stable for TES than for TSS, whereas TSS differences were smaller and more chemistry-dependent.

For TSS, annotator-dependent variance was most pronounced in CapTrap-enriched libraries, with particularly strong separation in PacBio-CapTrap (P = 1.28×10LL; ANOVA; **Supplementary Table S4B**) and a similarly clear effect in ONT-CapTrap (P = 2.94×10LL; **Supplementary Table S4B**). In both CapTrap datasets, LoRTIA Plus achieved the strongest mean TSS accuracy and retained significant advantages over all four competing annotators after BH-FDR correction **(Supplementary Table S4C)**. Relative to the best competing method, typical CapTrap TSS shifts were on the order of ∼0.06 F1 (e.g., ONT-CapTrap LoRTIA Plus vs bambu ΔF1 = 0.065; PacBio-CapTrap LoRTIA Plus vs FLAIR ΔF1 = 0.058), and the direction of the effect was concordant across all three cell lines. Annotator effects remained detectable in ONT-cDNA (P = 3.48×10LL) and PacBio cDNA (P = 8.70×10L³) as well, but differences were smaller and more heterogeneous. In ONT-cDNA, bambu and IsoQuant formed the top tier on mean TSS F1 **(Supplementary Table S4A)**, while LoRTIA Plus showed high recall that preserved a significant advantage over FLAIR (ΔF1 = 0.020; BH-adjusted P = 0.0060; **Supplementary Table S4C**) but did not exceed bambu or IsoQuant. In PacBio cDNA, LoRTIA Plus and FLAIR yielded closely similar mean TSS performance, consistent with reduced effect sizes outside CapTrap enrichment **(Supplementary Tables S4A–S4C)**.

For TES, ANOVA indicated strong annotator effects across all chemistries tested, including ONT-dRNA **(Supplementary Table S5B)**. In PCR-based libraries, LoRTIA Plus significantly outperformed each competing annotator in every chemistry after BH-FDR correction **(Supplementary Table S5C)**. Effect sizes were particularly large relative to NAGATA (ΔF1 ≈ 0.44–0.52 depending on chemistry), while contrasts versus bambu/IsoQuant/FLAIR were smaller but generally remained clear and statistically supported **(Supplementary Table S5C)**. Importantly, LoRTIA Plus’s TES advantage was not confined to CapTrap but recurred across the full set of PCR chemistries.

In ONT-dRNA libraries, TES ranking remained LoRTIA Plus > FLAIR > (bambu/IsoQuant) > NAGATA, with LoRTIA Plus retaining significant advantages over each competing annotator after correction **(Supplementary Table S5C)**. Notably, the separation between LoRTIA Plus and FLAIR, while statistically supported in this benchmark (ΔF1 = 0.140; BH-adjusted P = 0.015), was smaller than LoRTIA Plus’s contrasts versus IsoQuant, bambu and especially NAGATA, consistent with the expectation that reference-guided and de novo approaches can converge in TES accuracy on dRNA data for catalog-supported transcripts while differing substantially in how they explore the novel isoform space.

The dRNA-specific TES behavior of FLAIR and LoRTIA Plus likely reflects their design principles. FLAIR is reference-guided, anchoring long reads to the GENCODE transcript catalog prior to calling novel isoforms, whereas LoRTIA Plus operates fully de novo on genome-aligned reads and applies poly(A)-motif–based boundary detection specifically at the 3′ ends of ONT-dRNA reads. Accordingly, TES performance can appear comparable for annotated transcripts in this benchmark, while diverging sharply for novel NNC isoforms. When restricting the analysis to novel NNC isoforms—those lacking full or incomplete splice matches to the catalog and containing at least one unannotated donor or acceptor site—the contrast became extreme: FLAIR reported only a few dozen TES per dRNA sample with essentially negligible accuracy (TES F1 ≈ 0.001), whereas LoRTIA Plus detected several thousand novel TES per sample with high precision (≥0.9) and consistent support from an independent multi-chemistry + PolyASite reference set. Importantly, for ONT-dRNA novel TES, the benchmarked “false positive” rate can be inflated by catalog incompleteness. Under a catalog-based benchmark, dRNA-supported novel ends absent from GENCODE are counted as false positives, regardless of biological validity. This bias favors reference-guided approaches, which typically do not call (or call only sparsely) 3′ ends absent from the catalog, while de novo callers will recover them. The SPARC locus illustrates this directly **(Supplementary Figure S2)**: LoRTIA Plus identifies two dRNA-supported novel TES that are absent from both GENCODE and the FLAIR output, despite clear dRNA coverage supporting their existence. Conversely, a more distal GENCODE isoform with an extended 3′ UTR shows no detectable dRNA coverage and is not recovered by FLAIR. Thus, the apparent “false positive” inflation observed in the benchmark does not necessarily reflect erroneous calls, but often reflects gaps in the reference catalog.

In summary, boundary-level ANOVA and matched paired comparisons demonstrate that endpoint accuracy depends on both chemistry and algorithmic strategy. For TSS, the strongest and most reproducible method-dependent separation occurs in CapTrap-enriched protocols—most prominently in PacBio-CapTrap—where LoRTIA Plus provides the highest accuracy across all three cell lines. For TES, LoRTIA Plus provides the most accurate and most consistent performance across all evaluated chemistries, including ONT-dRNA, yielding a stable top rank with substantial effect sizes relative to all competitors **(Figure 3; Supplementary Tables S4–S5)**. Together, these results indicate that LoRTIA Plus not only matches or exceeds alternative methods on catalog-supported transcript boundaries, but also provides a robust framework for recovering reliable transcription start and end sites across diverse long-read library preparations.

### Catalog-supported splice-architecture recovery (SQANTI FSM/ISM)

We next evaluated recovery of known, GENCODE-supported splice architectures using SQANTI3 full-splice match (FSM) and incomplete-splice match (ISM) classifications. Because FSM and ISM are defined by concordance of exon–intron structure with annotated GENCODE reference transcripts, this analysis should be interpreted primarily as a junction-/intron-level evaluation of known transcript reconstruction; transcript-boundary accuracy was assessed separately in the preceding section. Accordingly, we quantified (i) FSM+ISM recovery, defined as the fraction of reference transcripts recovered as FSM or ISM, and (ii) the proportion of recovered transcripts classified as FSM [FSM/(FSM+ISM)].

Annotator choice produced robust and chemistry-dependent differences in both FSM+ISM recovery and the fraction of recovered transcripts classified as FSM [FSM/(FSM+ISM)]. For FSM+ISM recovery, chemistry-resolved ANOVA indicated strong annotator effects across all library types (ONT-CapTrap P = 5.0×10L¹¹; ONT-cDNA P = 9.1×10LL; ONT-dRNA P = 3.0×10LL; PacBio cDNA P = 9.0×10LL; PacBio-CapTrap P = 3.6×10LL; **Supplementary Table S6B**), whereas cell line had no significant effect in these models (P ≈ 0.21–0.96; **Supplementary Table S6B**), consistent with stable method ranking across the three matched cell lines. In contrast, for the fraction of recovered transcripts classified as FSM rather than ISM, annotator effects remained significant in all chemistries (P = 4.5×10L¹² to 7.7×10LL; **Supplementary Table S6B**) and cell-line effects were also detectable (P = 5.4×10LL to 0.0106; **Supplementary Table S6B**), suggesting that the balance between FSM and ISM reconstructions is more sensitive to biological and coverage differences.

Across all chemistries, LoRTIA Plus achieved the highest FSM+ISM recovery, i.e., it recovered the largest fraction of the GENCODE reference transcript set used in this benchmark **(Supplementary Table S6A)**. Averaged across the three cell lines, FSM+ISM recovery values (Recall_FSM_ISM) were 0.598 for ONT-CapTrap (second-best: FLAIR 0.519), 0.541 for ONT-cDNA (FLAIR 0.539; essentially tied), 0.594 for ONT-dRNA (bambu 0.580; close), 0.719 for PacBio cDNA (bambu 0.507), and 0.654 for PacBio-CapTrap (bambu 0.505) **(Supplementary Table S6A)**. Collapsing across all chemistry × cell-line conditions, LoRTIA Plus retained the top overall FSM+ISM recovery (0.621), followed by bambu (0.495), FLAIR (0.447), IsoQuant (0.274), and NAGATA (0.155) **(Supplementary Table S6A)**. Pre-specified paired comparisons (LoRTIA Plus versus each competitor, paired over cell lines with BH-FDR correction within each chemistry) confirmed large and significant FSM+ISM recovery gains over IsoQuant in every chemistry (e.g., PacBio Δ = +0.419, BH-adjusted P = 8.9×10LL; **Supplementary Table S6C)** and typically also over FLAIR, with ONT-cDNA as the exception where LoRTIA Plus and FLAIR did not differ (Δ = +0.0017,

BH-adjusted P = 0.88; **Supplementary Table S6C**). Comparisons against bambu were chemistry-dependent: LoRTIA Plus separated clearly in several CapTrap- and PacBio-derived datasets but showed only small, non-significant differences in ONT-cDNA and ONT-dRNA, consistent with the close mean FSM+ISM recovery values observed in those settings **(Supplementary Table S6C)**. These trends are summarized as chemistry-resolved effect sizes in **Figure 4A**.

**Figure 4.**
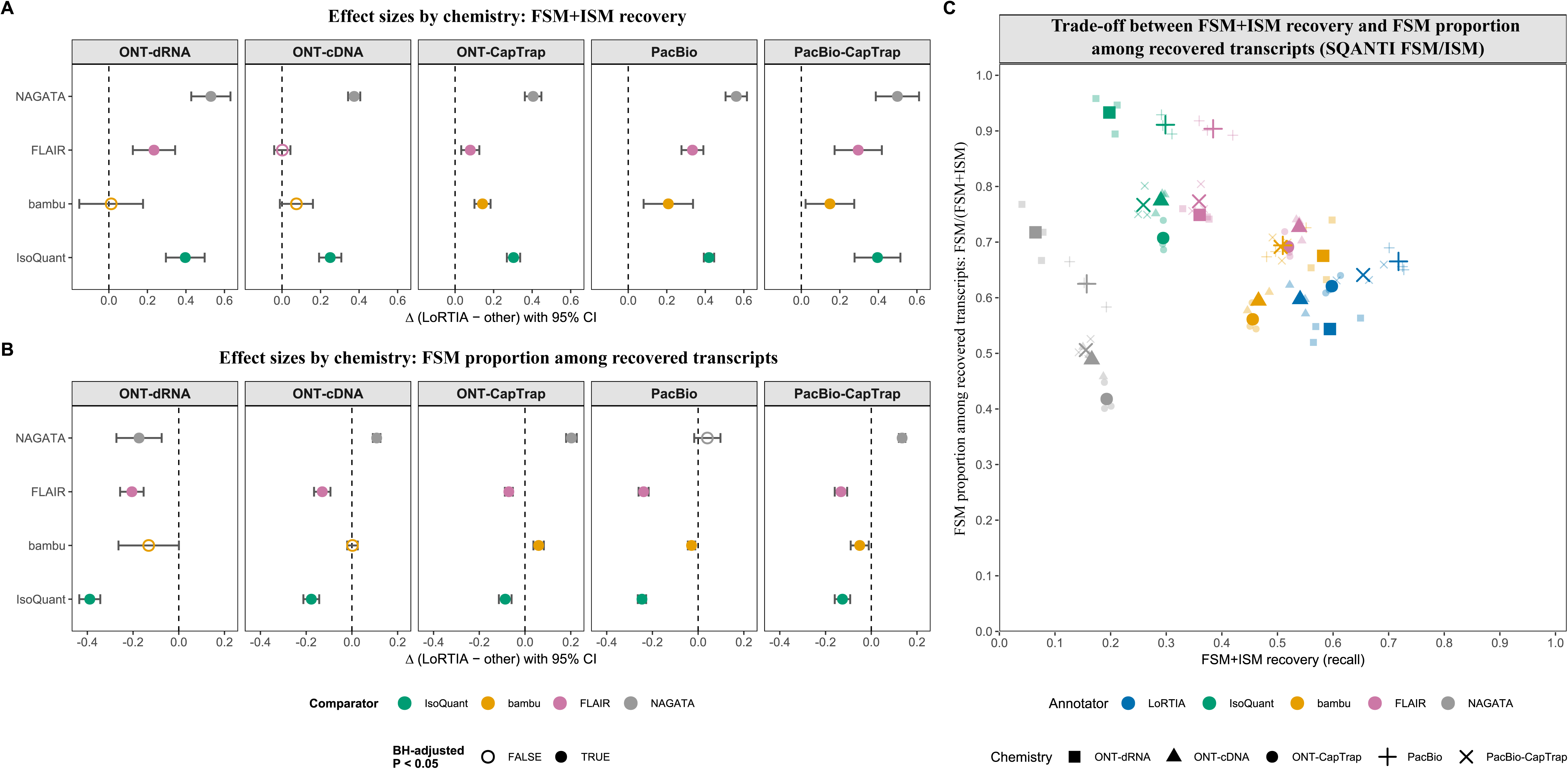
Transcript-level splice-architecture performance across annotators and long-read chemistries (SQANTI FSM/ISM). **(A)** Chemistry-resolved effect sizes for FSM+ISM recovery on known reference transcripts, quantified as the fraction of reference transcripts recovered as FSM or ISM in SQANTI3. Points show the mean paired difference Δ = FSM+ISM recovery {LoRTIA Plus} – FSM+ISM recovery_{other} across the three matched cell lines, with 95% confidence intervals from cell-line–paired differences; filled symbols denote BH-FDR–significant contrasts (adjusted *P* < 0.05) within each chemistry. **(B)** Chemistry-resolved effect sizes for the proportion of recovered transcripts classified as FSM rather than ISM, quantified as FSM/(FSM+ISM). Points and confidence intervals are computed as in panel A for Δ = [FSM/(FSM+ISM)]_{LoRTIA Plus} − [FSM/(FSM+ISM)]_{other}, where positive values indicate a higher FSM proportion among recovered transcripts. **(C)** Trade-off between FSM+ISM recovery and the proportion of recovered transcripts classified as FSM rather than ISM across annotators and chemistries. Each point represents an annotator × chemistry × cell-line combination; large markers indicate chemistry-specific centroids (means across cell lines). FSM+ISM recovery is plotted against FSM/(FSM+ISM) to visualize the method-dependent balance between broader recovery of known reference transcripts and a higher FSM proportion among recovered transcripts. Alt text: Comparison of splice-architecture performance across annotators and chemistries, showing recovery of known reference transcripts and the balance between FSM and ISM assignments.

However, the proportion of recovered transcripts classified as FSM [FSM/(FSM+ISM)] generally favored reference-guided approaches, particularly IsoQuant and often FLAIR, which showed higher FSM proportions among recovered transcripts **(Supplementary Table S6A)**. For example, in ONT-dRNA data IsoQuant and FLAIR reached ∼0.93 and ∼0.88, respectively, whereas LoRTIA Plus was ∼0.55; in PacBio cDNA, IsoQuant and FLAIR were ∼0.91 and ∼0.90 versus LoRTIA Plus at ∼0.66; and in ONT-CapTrap, IsoQuant/FLAIR were ∼0.85 versus LoRTIA Plus at ∼0.69 **(Supplementary Table S6A)**. Paired post hoc tests indicated that LoRTIA Plus exhibited significantly lower FSM proportions among recovered transcripts [FSM/(FSM+ISM)] than IsoQuant and FLAIR across chemistries (negative Δ with BH-adjusted P < 0.01; **Supplementary Table S6C**), whereas contrasts with bambu were mixed and chemistry-dependent **(Supplementary Table S6C)**. Chemistry-resolved effect sizes for the proportion of recovered transcripts classified as FSM [FSM/(FSM+ISM)] are shown in **Figure 4B**.

Together, the SQANTI FSM/ISM results reveal a clear and interpretable trade-off in splice-driven structural concordance on catalog-supported transcripts: LoRTIA Plus maximizes FSM+ISM recovery of known reference transcripts across long-read chemistries, while reference-guided pipelines increase the proportion of recovered transcripts classified as FSM rather than ISM. This trade-off between breadth of FSM+ISM recovery and FSM proportion among recovered transcripts is visualized directly by the joint distribution of the two metrics in **Figure 4C**. Because FSM/ISM classifications do not capture boundary accuracy, these splice-architecture results should be interpreted in conjunction with the dedicated TSS/TES endpoint benchmarking presented in the preceding section **(Figure 3)**; taken together, the two analyses provide a complementary view of method performance along the structural (junction) and boundary (start/end) dimensions.

Overall, LoRTIA Plus provides the most comprehensive splice-consistent recovery of known reference transcripts across chemistries **(Figure 4A)**, while IsoQuant and FLAIR show higher FSM proportions among recovered isoforms **(Figure 4B)**. The combined view **(Figure 4C)** highlights that method choice induces a consistent and chemistry-dependent shift between breadth of recovery and FSM proportion among recovered transcripts, supporting LoRTIA Plus as a competitive de novo annotator whose primary advantage lies in breadth of FSM+ISM recovery rather than maximal FSM fraction.

### Novel transcript discovery (NIC/NNC)

Because NIC and NNC categories represent transcript novelty outside the reference catalog rather than recovery of known models, we evaluated them not by reference-based accuracy, but by abundance, reproducibility and structural plausibility. We therefore examined novel transcript discovery along three complementary dimensions: (i) the abundance of NIC and NNC isoforms, (ii) reproducibility of novel splice junctions across annotators, chemistries and cell lines, and (iii) structural plausibility of novel junctions and isoforms **(Figure 5; Supplementary Table S7)**.

**Figure 5.**
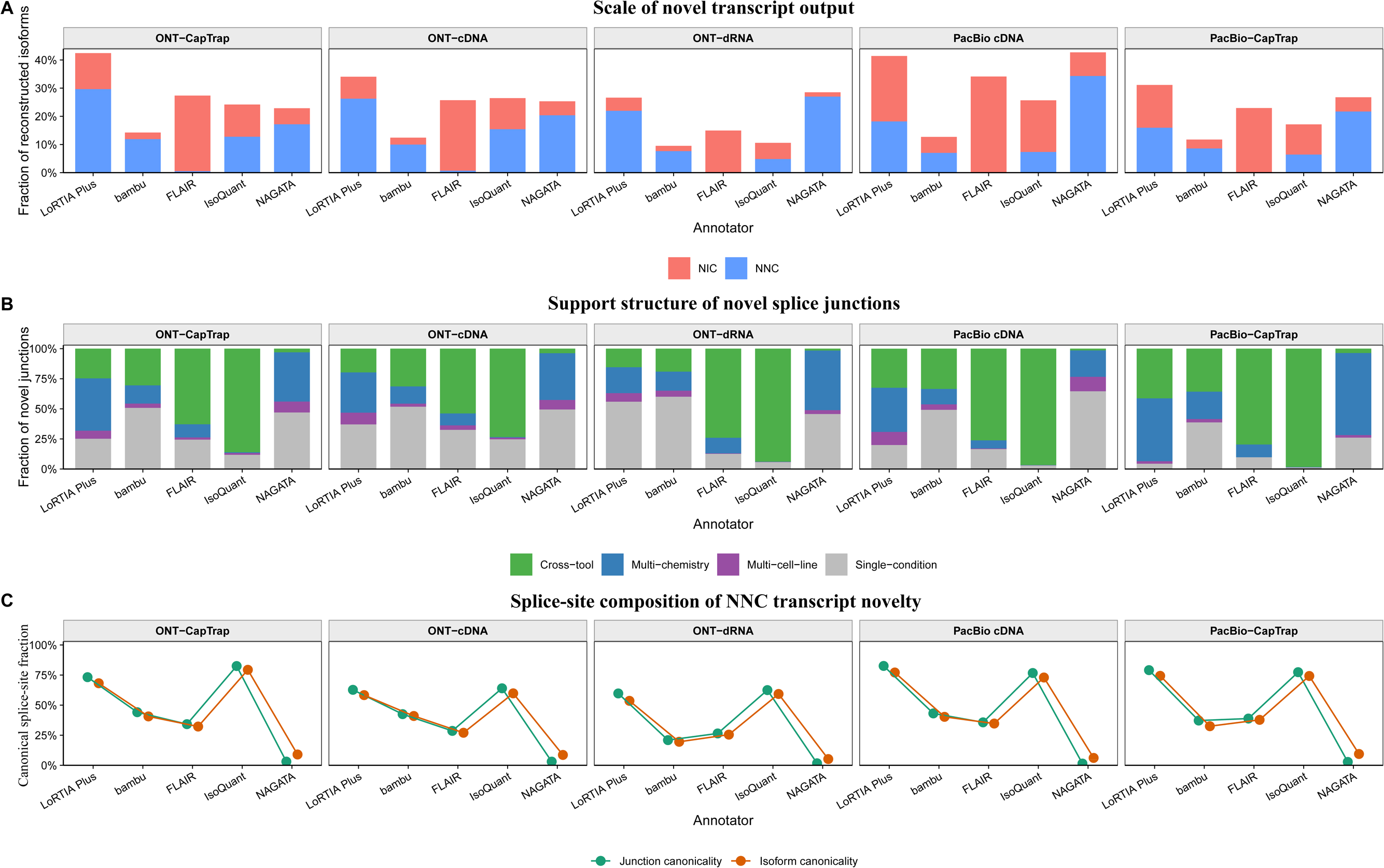
Novel transcript discovery and support structure across annotators and long-read chemistries. (A) Scale of novel transcript output across annotators and long-read chemistries, shown as the fraction of reconstructed isoforms assigned to the SQANTI3 categories novel in catalog (NIC) and novel not in catalog (NNC). Bars are partitioned into NIC and NNC components to illustrate both the scale and composition of novel transcript output. (B) Support structure of novel splice junctions across annotators and chemistries. Novel junctions were classified according to whether the same genomic donor–acceptor pair was detected by multiple annotators (cross-tool), recurred across multiple chemistries within one annotator (multi-chemistry), recurred across multiple cell lines within one annotator (multi-cell-line), or were restricted to a single annotator × chemistry × cell-line condition (single-condition). Bars show the fraction of novel junctions in each support class. (C) Canonical splice-site support in NNC transcript novelty. For each annotator and chemistry, the panel shows the fraction of novel NNC junctions carrying canonical donor–acceptor splice-site motifs (GT–AG, GC–AG or AT–AC; junction canonicality) together with the fraction of NNC isoforms carrying intron chains composed entirely of such junctions (isoform canonicality). This panel summarizes whether NNC transcript novelty is supported primarily by canonical splice-site architectures. Alt text: Comparison of novel transcript discovery across annotators and chemistries, including NIC/NNC output, support structure of novel junctions, and canonical splice-site support in novel isoforms.

Annotators differed markedly in both the scale and composition of novel transcript output. LoRTIA Plus annotated the most novel isoforms, with substantial NIC and NNC components. By contrast, FLAIR showed a predominantly NIC-skewed profile with minimal NNC output, indicating that it primarily reconstructs new combinations of previously annotated splice sites. NAGATA displayed an even more strongly NNC-enriched profile, whereas IsoQuant occupied an intermediate position and bambu remained the most conservative overall **(Figure 5A; Supplementary Table S7A)**. Thus, annotator choice shaped not only how much novelty was reported, but also the balance between catalog-compatible and more structurally novel transcript classes.

We next asked whether this novelty was independently supported. Novel splice junctions were classified according to whether the same donor–acceptor pair was detected by multiple annotators, recurred across multiple chemistries or cell lines, or remained restricted to a single annotator × chemistry × cell-line condition. In this analysis, a substantial fraction of the novel junctions reported by LoRTIA Plus showed independent support, indicating that its novel splice-junction output was not dominated by single events. By contrast, although NAGATA expanded the set of novel junctions more strongly, a much larger fraction of these calls remained condition-specific, consistent with weaker reproducibility. IsoQuant and FLAIR covered smaller sets of novel junctions, but the novelty they reported showed higher relative support across independent contexts, in line with their more reference-constrained behavior. bambu showed an intermediate pattern, retaining a measurable discovery component but with fewer novel junctions than LoRTIA Plus **(Figure 5B; Supplementary Table S7B)**. These results indicate that a large component of the novel splice-junction output recovered by LoRTIA Plus is reproducible across independent analytical or experimental contexts.

We then evaluated structural plausibility using splice-site composition. At the junction level, LoRTIA Plus showed a high fraction of canonical donor–acceptor splice-site motifs (GT–AG, GC–AG or AT–AC) among novel junctions, and this remained true in the more stringent NNC-only subset, i.e. among events containing at least one previously unannotated donor or acceptor site. In this view, many NNC isoforms recovered by LoRTIA Plus were composed of junctions with canonical splice-site motifs, broadly comparable to IsoQuant but substantially larger in scale. FLAIR also showed a high fraction of novel junctions carrying canonical splice-site motifs across its full novel junction set, but this primarily reflected its NIC-dominant profile and its limited NNC output. In contrast, NAGATA recovered a large number of NNC isoforms, but most of this novelty was associated with non-canonical splice-site motifs, indicating substantially weaker structural plausibility **(Figure 5C; Supplementary Table S7C)**. Isoform-level summaries were concordant with these junction-level results, showing that a substantial fraction of both NIC and NNC isoforms reported by LoRTIA Plus carried fully canonical intron chains, whereas NAGATA retained a much larger non-canonical component **(Supplementary Table S7D)**. Thus, the novel transcript output reported by LoRTIA Plus is not only extensive, but also structurally credible.

Overall, novel transcript discovery could not be interpreted from novelty counts alone. Although NAGATA expanded the set of novel junctions and novel isoforms more aggressively, much of that output remained condition-specific and structurally less convincing. LoRTIA Plus, by contrast, combined a large novel transcript output with substantial independent support and strong structural plausibility. This pattern argues that the advantage of LoRTIA Plus in novel transcript discovery does not arise merely from permissive overcalling, but from recovery of a larger and more biologically credible set of de novo transcript models.

## Discussion

Across viral and human benchmarks, LoRTIA Plus delivered robust, high-fidelity transcript annotation that generalized across long-read sequencing chemistries. In KSHV, where an orthogonally supported reference set of TSSs and TESs was available, LoRTIA Plus achieved the strongest performance for transcript-boundary recovery in both direct cDNA and direct RNA datasets, and this advantage extended to transcript-level reconstruction. In the human LRGASP datasets spanning three cell types and five long-read chemistries, LoRTIA Plus consistently ranked among the strongest annotators in boundary-focused benchmarking and, particularly in TES benchmarking and CapTrap-enriched TSS analyses, often emerged as the most accurate method while remaining competitive in ONT dRNA data. Taken together, these results support the central premise of LoRTIA Plus: a chemistry-agnostic, feature-first strategy—in which TSSs, TESs and introns are validated and filtered before transcript assembly—can improve endpoint fidelity without sacrificing the discovery of biologically meaningful novel isoforms.

The KSHV benchmark illustrates particularly clearly why explicit treatment of transcript boundaries is essential in compact, gene-dense genomes. Overlapping coding regions, convergent and divergent promoter architectures, and pervasive readthrough transcription create conditions in which splice-identical but boundary-distinct isoforms can easily collapse into a single “canonical” transcript, masking regulatory diversity encoded at the level of alternative 5′ and 3′ ends. Methods that rely primarily on splice junctions while treating transcript ends as secondary features are especially vulnerable to such boundary collapse (6, 12, 13). By contrast, LoRTIA Plus treats TSSs, TESs and introns as primary entities and assembles transcript models only from reads supported at both ends and across all retained splice junctions. Adapter-aware boundary quality control further reduces internal priming and template-switching artifacts and enriches for molecules with intact 5′ ends in cDNA libraries. In the KSHV system, this design translated into consistently superior TSS-, TES- and transcript-level performance across both ONT chemistries, indicating that feature-first, adapter-aware annotation is particularly well suited to dense viral transcriptomes.

The human LRGASP analyses further showed that annotator choice shapes the observed transcriptome at least as strongly as library chemistry. SQANTI3 structural classes, transcript length distributions, coding/non-coding balance and gene-level isoform complexity collectively revealed distinct and reproducible annotation regimes. IsoQuant and FLAIR represented more reference-constrained profiles, with higher FSM fractions and more constrained novel transcript output, whereas NAGATA occupied a more permissive regime characterized by low FSM fractions, abundant antisense and NNC output, and more fragmented transcript structures. bambu lay between these extremes, producing largely splice-consistent but ISM-enriched models. LoRTIA Plus combined strong recovery of known transcript structures with broader novel transcript output. In several global structural metrics, between-tool differences were comparable to, or greater than, between-chemistry differences, emphasizing that long-read transcriptomics does not yield a single tool-independent view of transcriptome structure but rather method-dependent representations of the same underlying RNA population.

Boundary-focused benchmarking clarified more precisely where LoRTIA Plus sits within this landscape. For both TSS and TES detection, method effects were strongest in libraries with improved end representation, especially CapTrap-based protocols. Across the three LRGASP cell lines, LoRTIA Plus provided the most accurate TSS and, in particular, the most consistent TES calls in CapTrap libraries. Its TES advantage was not confined to CapTrap, but extended across the broader set of PCR-based chemistries. Because CapTrap-like protocols improve representation of transcript termini, these results suggest that high-end-fidelity library preparation combined with feature-level validation is especially effective for workflows prioritizing precise transcript boundary definition and reference refinement (31).

At the same time, splice-architecture recovery and endpoint fidelity measure distinct aspects of annotation performance. This distinction was reflected in the SQANTI FSM/ISM analysis: LoRTIA Plus did not maximize the proportion of recovered transcripts classified as FSM rather than ISM, but instead achieved the highest FSM+ISM recovery. In other words, it recovered the largest fraction of known, GENCODE-supported transcript models, whereas reference-guided methods—particularly IsoQuant and, in many chemistries, FLAIR—produced higher FSM proportions among the transcripts they recovered. This reveals a clear and interpretable trade-off. LoRTIA Plus is strongest in broad recovery of splice-consistent known transcript structures, whereas reference-guided pipelines tend to maximize the fraction of FSM classifications among recovered isoforms. The resulting trade-off between breadth of FSM+ISM recovery and FSM proportion among recovered transcripts indicates that splice-architecture benchmarking is not reducible to a simple ranking of tools from “better” to “worse”, but depends on whether the analytical priority is maximal recovery of known models or a higher FSM fraction among recovered isoforms.

One of the most important findings of the current study is that novel transcript discovery cannot be evaluated by NIC/NNC counts alone. LoRTIA Plus indeed produced the broadest novel transcript output and did so with a substantial NNC component. However, support structure and splice-site composition both indicated that this discovery output was not dominated by random or private events. A substantial fraction of novel junctions reported by LoRTIA Plus showed independent support, either through recurrence across annotators or through recurrence across chemistries and cell lines. Moreover, much of the NNC output was supported by canonical donor–acceptor motifs at the junction level and by exon chains composed entirely of such junctions at the isoform level. By contrast, NAGATA expanded novel transcript output even more aggressively, but a much larger fraction of that output remained condition-specific and was associated with non-consensus splice-site patterns. IsoQuant and FLAIR produced smaller but more constrained and highly supported novel outputs. Together, these results suggest that the advantage of LoRTIA Plus in novel transcript discovery does not arise merely from permissive overcalling, but from recovery of a broader, reproducible and structurally plausible set of de novo transcript models.

ONT dRNA data require separate interpretation. In this chemistry, threading of RNA into the motor–pore complex typically results in loss of the extreme 5′ terminal nucleotides, while basecalling uncertainty near the poly(A) tail complicates endpoint identification at the 3′ end (32–34). These chemistry-specific constraints help explain why reference-guided approaches may appear competitive in some TES comparisons restricted to known transcript structures, even when they underperform in the genuinely novel transcript space. Our results are fully consistent with this view: in ONT dRNA, the strongest advantage of LoRTIA Plus emerged in the novel TES space, where de novo end detection combined with poly(A)-motif-based filtering captured transcript ends that are unlikely to be recovered by reference-anchored pipelines. Looking forward, dRNA protocols that improve 5′ cap preservation or recover missing 5′ information through adapter ligation are likely to further increase the utility of feature-first, adapter-aware boundary detection in human transcriptomics as well.

These findings have practical implications. In long-read transcriptome annotation, library chemistry and annotator choice should not be treated as independent variables. Protocols that preserve or enrich transcript termini—such as CapTrap or well-optimized cDNA workflows—are especially well matched to methods such as LoRTIA Plus that explicitly validate and filter transcript features before model assembly. At the same time, our analyses caution against interpreting any single transcript annotation as definitive. Conservative approaches may maximize agreement with the existing reference catalog but miss alternative UTR structures and rarer isoforms, whereas permissive approaches may greatly expand the novel transcript space at the cost of increased noise. LoRTIA Plus occupies an intermediate position that is likely to be advantageous in studies where both recovery of known transcript structures and reliable novel discovery are priorities.

This study also has limitations. The KSHV reference set, although assembled from multiple assays and publications, is necessarily incomplete; low-abundance isoforms and rare boundary events are likely to remain unannotated and may therefore appear as apparent false positives. In human data, the TSS and TES truth sets rely on CAGE-, PolyASite- and GENCODE/SQANTI3-based resources, each of which has its own detection limits and positional uncertainty. In addition, the human LRGASP and KSHV benchmark frameworks were intentionally restricted to reference features recovered by at least one annotator within the defined tolerance. This design reduces the influence of biologically inactive or practically undetectable reference features, but also focuses evaluation on the accessible reference space rather than the complete reference set. This can bias benchmarking toward underestimation of true performance: biologically real transcript ends or isoforms absent from these resources will be scored as apparent false positives, and small coordinate differences can convert near-matches into formal mismatches. In addition, although we evaluated a representative set of annotators, further chemistry-aware parameter optimization—particularly of support thresholds, artifact filters and endpoint tolerances—could likely improve individual pipelines. Despite these caveats, our results position LoRTIA Plus as a chemistry-agnostic solution for high-fidelity transcriptome reconstruction that does not depend on pre-existing transcript annotations. By combining adapter-aware boundary quality control, motif- and statistics-based feature validation, and a filter-first, assemble-later design, LoRTIA Plus simultaneously preserves strong reference concordance, improves recovery of true transcript boundaries and splice architectures, and expands the space of reproducible novel transcript models.

In conclusion, LoRTIA Plus provides a practical, chemistry-agnostic framework for high-fidelity transcriptome annotation that complements existing methods by prioritizing both endpoint fidelity and biologically credible novel isoform discovery. Its open-source implementation, detailed documentation and compatibility with current ONT and PacBio platforms position it as a valuable addition to the long-read transcriptomics toolkit, particularly for studies requiring precise transcript-boundary delineation together with robust recovery of novel transcript diversity.

## Data availability

KSHV long-read datasets (ONT dRNA-seq and dcDNA-seq) are available under GEO Series accessions GSE250427 and GSE250429, and under ENA BioProject accession PRJEB60022. Human LRGASP datasets are available via the ENCODE Portal under the following experiment accessions: H1-hES: ENCSR016IKV (ONT cDNA), ENCSR049AYR (ONT dRNA), ENCSR522NAJ (ONT CapTrap), ENCSR271KEJ (PacBio cDNA); H1-DE: ENCSR485VRY (ONT cDNA), ENCSR758ABF (ONT dRNA), ENCSR543ORO (ONT CapTrap), ENCSR127HKN (PacBio cDNA); WTC11: ENCSR539ZXJ (ONT cDNA), ENCSR392BGY (ONT dRNA), ENCSR054ABL (ONT CapTrap), ENCSR507JOF (PacBio cDNA), ENCSR309IKK (PacBio CapTrap).

## Author contributions

G.T.: Conceptualization, benchmarking design, formal analysis, software development, visualization, writing—original draft. Z.Ba.: Conceptualization, software development, writing—review and editing. D.T.: Conceptualization, supervision, writing—review and editing. Z.Bo.: Conceptualization, supervision, project administration, writing—review and editing. Á.F.: Data curation, supporting analyses. All authors approved the final version of the manuscript.

## Funding

National Research, Development and Innovation Office grants: K 142674 and ADVANCED 152705 to Zsolt Boldogkői. The APC was covered by the University of Szeged Open Access Fund: 8070.

## Declaration of usage of generative AI and AI-assisted technologies

No artificial intelligence–based tools were used for data processing, statistical analysis or interpretation of the results. AI assistance (ChatGPT, OpenAI) was used only for generating figure-plotting code and for language editing of the manuscript; all scientific content was conceived, checked and approved by the authors.

## Conflict of interest

The authors declare no competing interests.

## Notes

### Competing Interest Statement

The authors have declared no competing interest.

