## Supplementary Information for "LoRTIA Plus: a chemistry-agnostic, feature-first software package for long-read transcriptome annotation"

### Supplementary Figure S1. Cell line–resolved structural and complexity profiles across annotators and chemistries.

Extended view of the global structural properties, stratified by cell line (H1, H1-DE, WTC11). All panels show results for the five annotators (LoRTIA Plus, bambu, FLAIR, IsoQuant and NAGATA) across the five long-read chemistries (ONT-dRNA, ONT-cDNA, ONT-CapTrap, PacBio cDNA and PacBio-CapTrap), but separated by cell line rather than aggregated.

**(A)** SQANTI3 primary and secondary structural categories for each cell line. For every annotator–chemistry–cell-line combination, the proportions of transcripts assigned to full splice match (FSM), incomplete splice match (ISM), novel in catalog (NIC), novel not in catalog (NNC) and additional classes (Antisense, Genic Intron, Genic Genomic, Intergenic, Fusion) are shown, mirroring the aggregated profiles in Figure 2A but resolved separately for H1, H1-DE and WTC11.

**(B)** Transcript length distributions per cell line. For each annotator and chemistry, the distribution of isoform lengths is displayed separately for H1, H1-DE and WTC11, extending the global length summaries in Figure 2B. These panels confirm that the chemistry-driven differences in median length and the annotator-specific length patterns are highly reproducible across all three cell lines.

**(C)** Coding versus non-coding transcript fractions per cell line. For each annotator–chemistry–cell-line combination, the proportion of coding isoforms (those containing an intact ORF) versus non-coding models is shown, complementing the aggregated coding rates in Figure 2C. The per-cell-line profiles demonstrate that the relative ranking of annotators in terms of coding enrichment is stable across H1, H1-DE and WTC11.

**(D)** Number of isoforms per gene per cell line. For each annotator and chemistry, the distribution of isoform counts per gene is reported separately for H1, H1-DE and WTC11, extending the gene-level complexity summaries in Figure 2D. These panels show that the characteristic annotation patterns—compact models for FLAIR, intermediate complexity for bambu and IsoQuant, and richer isoform repertoires for LoRTIA Plus and NAGATA—are consistently observed in all three cell lines.

### Supplementary Figure S2. **SPARC locus illustrating dRNA-supported novel TES that are missed by reference-guided annotation.**

Genome-browser view of the SPARC locus on human chromosome 5, showing GENCODE reference transcripts (“Gencode”), the FLAIR reference-guided annotation (“FLAIR”) and LoRTIA Plus-derived isoforms (“LoRTIA Plus”). The label “Previously annotated TES” marks the canonical 3′ end that is present in GENCODE and is also recovered by FLAIR. In contrast, the two positions marked as “Novel TES” correspond to additional 3′ termini called only by LoRTIA Plus; these sites are not annotated in GENCODE and are therefore not detected by FLAIR, but are supported by ONT dRNA reads and local poly(A) motifs (see main text). A more distal GENCODE isoform with an extended 3′ UTR, located beyond the canonical TES, shows no detectable dRNA coverage in our data and is not reconstructed by FLAIR. Together, this example illustrates that many LoRTIA Plus-called TES that appear as “false positives” when evaluated solely against a static reference actually represent alternative, cell type–specific 3′ ends, whereas reference-guided approaches have limited ability to discover such events if they are not already present in the underlying catalog.

Supplementary Table S1. Benchmark datasets used for KSHV and human transcriptome analyses.

Benchmark datasets used for KSHV and human transcriptome analyses.

Supplementary Table S1A. KSHV benchmarking datasets and supporting reference resources.

This table compiles the sequencing datasets and supporting reference resources used in the KSHV and human benchmarking analyses. For the viral benchmark, it lists the long-read and complementary datasets used to define high-confidence reference transcription start sites, transcription end sites, and introns. For the human benchmark, it summarizes the LRGASP/ENCODE long-read datasets analysed across cell lines and sequencing chemistries, together with the corresponding accession identifiers. These resources form the basis of all downstream endpoint-, transcript-, and structure-level performance comparisons in this study.

### **Supplementary Table S1B. Human LRGASP/ENCODE long-read RNA-seq benchmark datasets.**

This table summarizes the human long-read RNA-seq datasets analysed in this study from the LRGASP/ENCODE resource, covering three cell lines (H1-hES, H1-DE, and WTC11) and five long-read library chemistries (ONT dRNA-seq, ONT cDNA-seq, ONT CapTrap-seq, PacBio cDNA-seq, and PacBio CapTrap-seq), together with the corresponding ENCODE accession identifiers.

Supplementary Table S2. KSHV reference transcript endpoints and benchmarking metrics.
This table compiles the assay- and literature-supported reference boundary sets used for KSHV benchmarking and reports performance metrics for LoRTIA Plus and alternative long-read transcript annotators across ONT dRNA (dRNA) and dcDNA (dcDNA) libraries (see Methods). High-confidence TSSs were restricted to literature-supported positions derived from published CAGE/RAMPAGE resources, while TESs were retained only when supported by poly(A)-anchored 3′-end evidence from ONT dRNA reads, providing an orthogonal reference for both ONT protocols.

Supplementary Table S2A. Endpoint-level benchmarking metrics (TSS and TES).
Precision, recall and F1 score for TSS and TES endpoints across the five annotators, reported separately for ONT dRNA and ONT dcDNA libraries under the matching criteria described in Methods.

Supplementary Table S2B. Transcript-level benchmarking metrics.
Transcript-level performance summaries for the five annotators on ONT dRNA and ONT dcDNA libraries, reported as transcript-level F1 scores under the joint TSS–TES correctness criterion defined in Methods.

Supplementary Table S2C. Transcript-class composition of true-positive (reference-matched) calls in ONT dcDNA.
Transcript-class distributions for **dcDNA predictions that matched the KSHV reference set** (i.e., true positives under the defined positional tolerance), summarized for each annotator.

Supplementary Table S2D. Transcript-class composition of true-positive (reference-matched) calls in ONT dRNA.
Transcript-class distributions for dRNA predictions that matched the KSHV reference set (true positives under the defined positional tolerance), summarized for each annotator.

Supplementary Table S2E. Transcript-class composition of false-positive (unmatched) calls in ONT dcDNA.
Transcript-class distributions for dcDNA predictions that did not match any reference transcript unit (false positives under the defined positional tolerance), summarized for each annotator.

Supplementary Table S2F. Transcript-class composition of false-positive (unmatched) calls in ONT dRNA.
Transcript-class distributions for dRNA predictions that did not match any reference transcript unit (false positives under the defined positional tolerance), summarized for each annotator.

### Supplementary Table S3. Global structural properties of human long-read transcriptomes across annotators and library chemistries.

This workbook provides the cell line–aggregated numerical summaries underlying Figure 2A–D, including SQANTI3 structural category composition, coding/non-coding balance, transcript length statistics, and gene-level isoform complexity across five long-read library chemistries and five transcript annotators.

Supplementary Table S3A. — SQANTI3 category and coding summary.
Chemistry-resolved, cell line–aggregated counts and fractions of SQANTI3 primary structural categories (FSM, ISM, NIC, NNC) and additional classes (e.g., antisense/genic/genic-intron/intergenic/fusion, as applicable), together with coding/non-coding fractions and related summary metrics used for Figure 2A and Figure 2C.

Supplementary Table S3B.— Transcript length distribution statistics.
Chemistry-resolved, cell line–aggregated transcript length summaries (e.g., median and interquartile range, with additional distribution descriptors where provided) for each annotator, corresponding to Figure 2B.

### Supplementary Table S3C. — Gene-level isoform complexity.

Chemistry-resolved, cell line–aggregated gene-level isoform complexity summaries (e.g., mean isoforms per gene and the fraction of genes exceeding specified isoform-count thresholds, where provided) for each annotator, corresponding to Figure 2D.

### Supplementary Table S3D. - Tool- and protocol-specific patterns of novel splice-junction classes

Novel splice-junction classes and their relative frequencies across chemistries, cell lines and annotators in human LRGASP datasets.

### Supplementary Table S4. Transcript boundary detection (TSS): accuracy metrics and statistical comparisons across annotators and library chemistries

This table summarizes transcription start site (TSS) boundary detection performance across annotators and long-read library chemistries using the three LRGASP cell lines as matched biological replicates.

Supplementary Table S4A. – TSS accuracy metrics (Precision/Recall/F1).
Boundary-level precision, recall, and F1 score for TSS detection for each annotator × chemistry × cell line combination. TSS benchmarking is reported for the four PCR-based chemistries (ONT-CapTrap, ONT-cDNA, PacBio cDNA, PacBio-CapTrap); ONT-dRNA is not evaluated for TSS due to frequent 5′-end truncation in direct RNA reads.

Supplementary Table S4B. – TSS repeated-measures ANOVA by chemistry.
Chemistry-resolved ANOVA testing the effect of Annotator on TSS F1 scores with cell line included as a blocking factor. Columns report the F statistic, degrees of freedom (df1, df2), and P value, along with the number of cell lines and annotators included.

Supplementary Table S4C. – TSS post hoc paired comparisons (LoRTIA Plus vs others).
Pre-specified cell-line–paired t-tests comparing LoRTIA Plus to each competing annotator within each chemistry. Reported are mean ΔF1 (LoRTIA Plus − other), SD of ΔF1, the t statistic, the two-sided P value, and the Benjamini–Hochberg (BH) FDR-adjusted P value (multiple-testing correction across the four contrasts within each chemistry).

### Supplementary Table S5. Transcript boundary detection (TES): accuracy metrics and statistical comparisons across annotators and library chemistries

This table summarizes transcription end site (TES) boundary detection performance across annotators and library chemistries, including ONT-dRNA, using the three LRGASP cell lines as matched biological replicates.

Supplementary Table S5A. – TES accuracy metrics (Precision/Recall/F1).
Boundary-level precision, recall, and F1 score for TES detection for each annotator × chemistry × cell line combination.

Supplementary Table S5B. – TES repeated-measures ANOVA by chemistry.
Chemistry-resolved ANOVA testing the effect of Annotator on TES F1 scores with cell line included as a blocking factor, reporting F, df1/df2, and P value.

Supplementary Table S5C. – TES post hoc paired comparisons (LoRTIA Plus vs others).
Pre-specified cell-line–paired t-tests comparing LoRTIA Plus to each competing annotator within each chemistry (including ONT-dRNA). Reported are mean ΔF1 (LoRTIA Plus − other), SD of ΔF1, the t statistic, the two-sided P value, and the Benjamini–Hochberg (BH) FDR-adjusted P value (multiple-testing correction across the four contrasts within each chemistry).

### Supplementary Table S6. Transcript-level splice-architecture benchmarking (SQANTI FSM/ISM) across annotators and long-read chemistries

This table summarizes SQANTI3-based transcript-level performance on known, GENCODE-supported transcript structures using FSM/ISM assignments across annotators and long-read library chemistries. Because FSM and ISM primarily reflect splice-architecture concordance with the GENCODE reference, the reported metrics quantify recovery of known transcript structures rather than transcript-boundary accuracy.

### Supplementary Table S6A. SQANTI transcript-level metrics (FSM/ISM)

FSM+ISM recovery and the proportion of recovered transcripts classified as FSM rather than ISM [FSM/(FSM+ISM)] for each annotator × chemistry × cell-line combination. Counts for the reference transcript set used in the benchmark and the recovered transcript set are also reported, together with FSM recovery and ISM recovery.

### Supplementary Table S6B. Chemistry-resolved ANOVA for SQANTI transcript-level metrics

ANOVA testing the effect of Annotator on (i) FSM+ISM recovery and (ii) the proportion of recovered transcripts classified as FSM rather than ISM [FSM/(FSM+ISM)], with cell line included in the model. Columns report F statistics, degrees of freedom and P values.

### Supplementary Table S6C. Post hoc paired comparisons (LoRTIA Plus vs other annotators)

Pre-specified cell-line–paired t-tests comparing LoRTIA Plus to each competing annotator within each chemistry, reported separately for FSM+ISM recovery and the proportion of recovered transcripts classified as FSM rather than ISM [FSM/(FSM+ISM)]. Shown are mean Δ, SD of Δ, Cohen’s dz, t statistic, raw P value and BH-FDR–adjusted P value.

### Supplementary Table S7. Novel transcript discovery and support structure across annotators and long-read chemistries

This table summarizes novel transcript discovery outside the GENCODE reference across the five annotators and five long-read chemistries in the three human LRGASP cell lines. The reported quantities describe the scale, reproducibility, and structural plausibility of novel transcript models and novel splice junctions, with separate summaries for NIC and NNC categories where appropriate.

### Supplementary Table S7A. Scale of novel transcript output (NIC/NNC)

Counts and fractions of novel in catalog (NIC) and novel not in catalog (NNC) transcript models for each annotator × chemistry × cell-line combination, together with annotator-level pooled summaries. Reported values include the total number of isoforms, NIC count, NNC count, total novel isoforms, the fraction of NIC and NNC isoforms among all reconstructed models, and the NNC/NIC ratio.

### Supplementary Table S7B. Support structure of novel splice junctions

Support-class summaries for novel splice junctions, defined by genomic donor–acceptor pairs, across annotators and chemistries. Novel junctions are classified according to whether they are shared across annotators, recur across multiple chemistries within one annotator, recur across multiple cell lines, or remain restricted to a single annotator × chemistry × cell-line condition. Reported values include the number of unique novel junctions and the fractions assigned to each support class, together with the overall fraction showing independent support.

### Supplementary Table S7C. Structural plausibility of novel splice junctions

Chemistry-resolved summaries of novel splice-junction composition across annotators, reported separately for all novel junctions and for the NNC-only subset. Reported values include the number of junction rows, the fraction of junctions carrying GT-AG, GC-AG or AT/AC donor–acceptor splice-site motifs, the corresponding fraction with non-consensus splice-site patterns, the relative contribution of GT-AG, GC-AG and AT-AC splice-site patterns, and the fraction of RTS-flagged junctions. The indel_near_fraction field was not informative in the current dataset and was therefore not interpreted further.

### Supplementary Table S7D. Isoform-level splice-site composition of NIC and NNC models

Chemistry-resolved isoform-level summaries for NIC and NNC transcript models. Reported values include the number of novel isoforms in each category, the fraction of isoforms carrying intron chains composed entirely of GT-AG, GC-AG or AT-AC junctions, the fraction of isoforms containing non-consensus splice-site patterns, the proportion of missing/undetermined splice-site composition assignments, predicted NMD fraction, bite fraction, and median transcript length and exon count.
