## Supplementary figures and images for "LoRTIA Plus: a chemistry-agnostic, feature-first software package for long-read transcriptome annotation"

### Supplementary Figure S1A

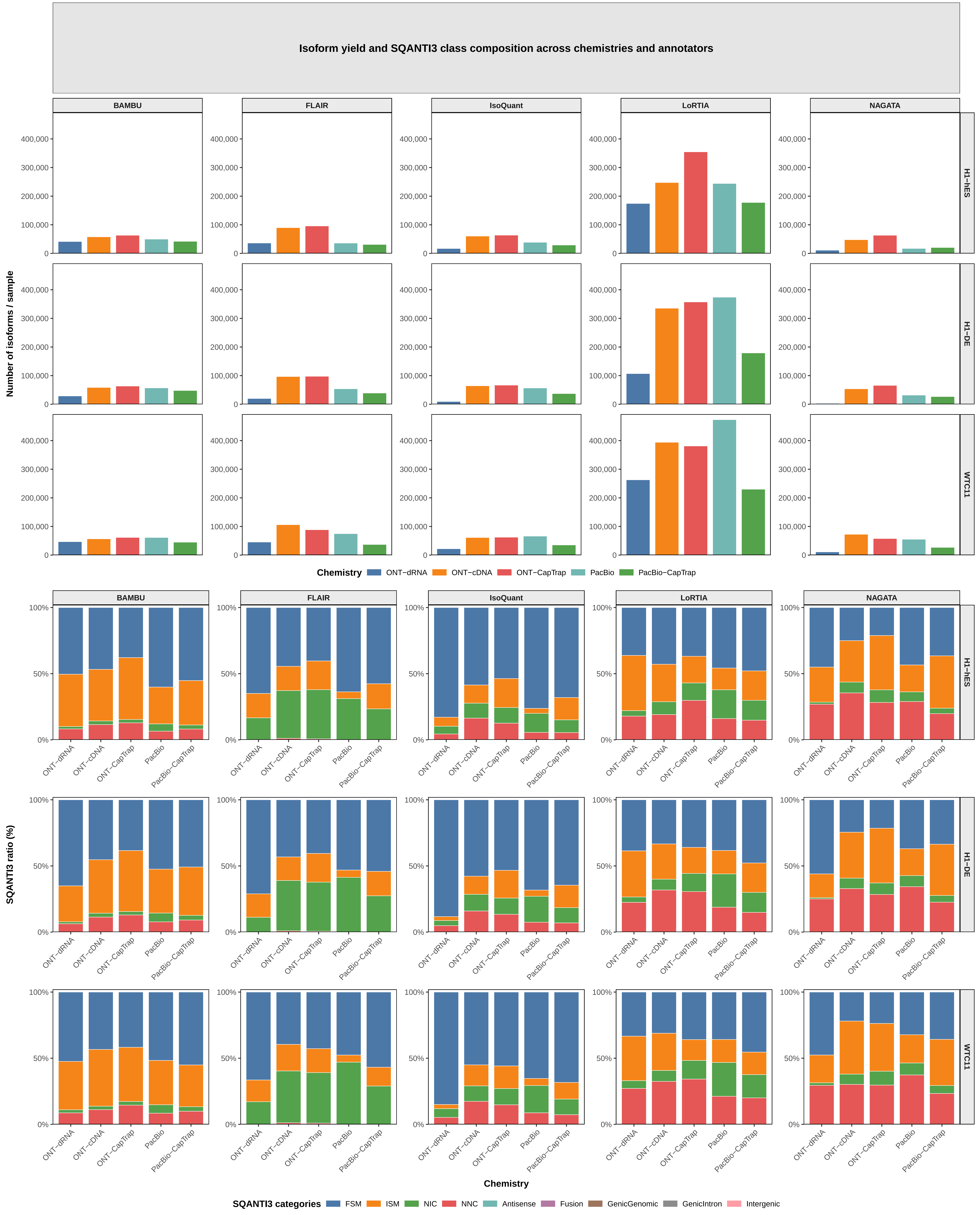

### Supplementary Figure S1B

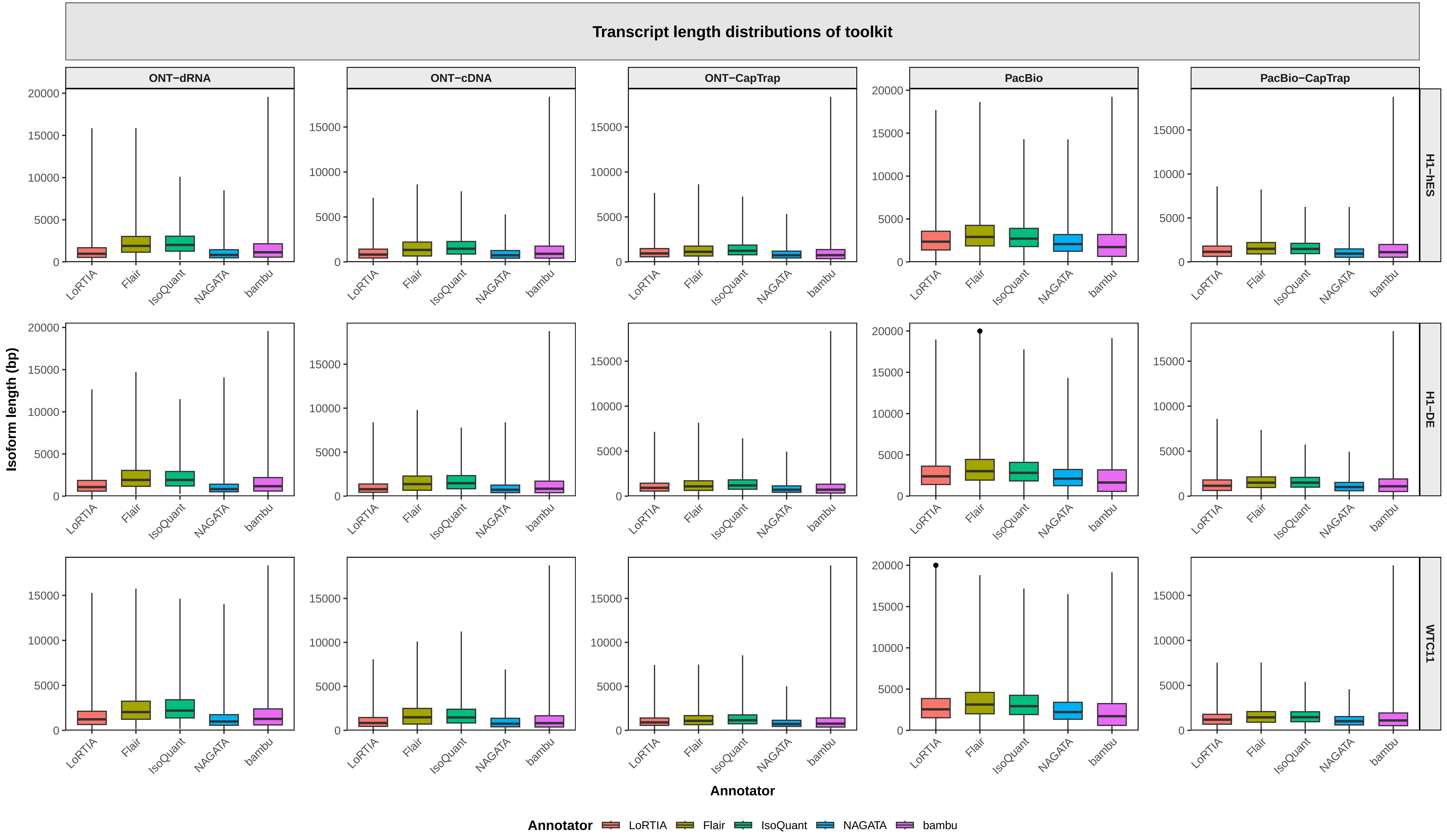

### Supplementary Figure S2

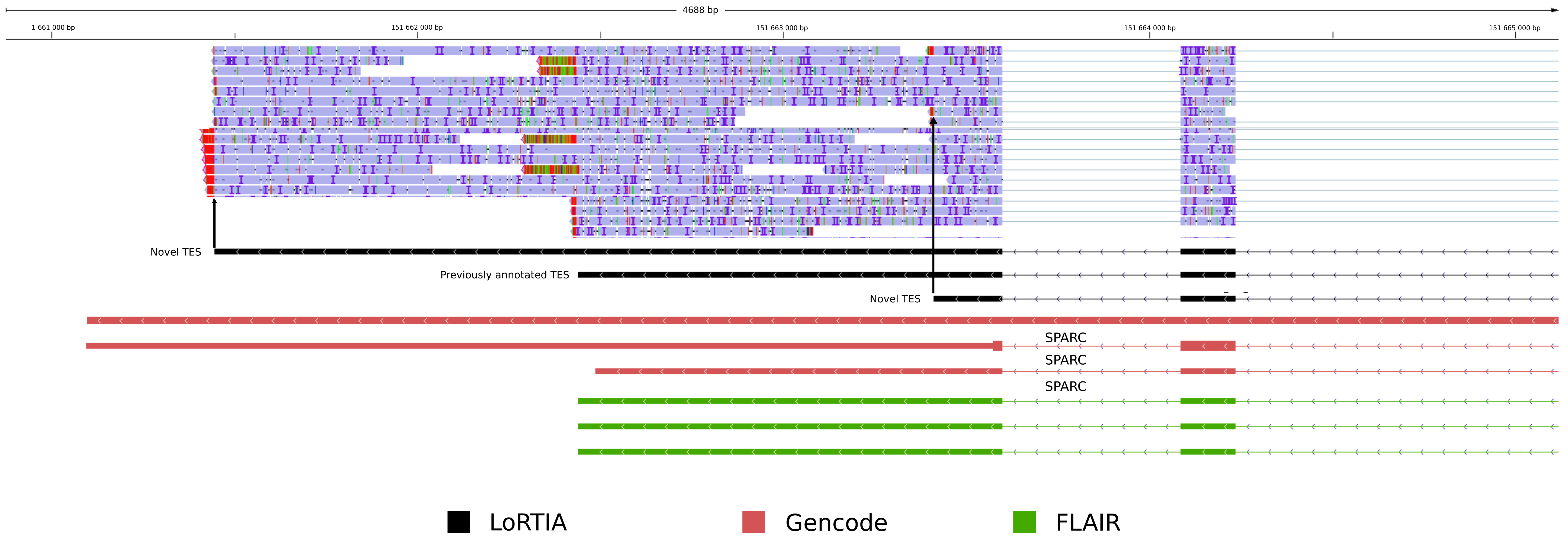
